# An Archaeal Transcription Factor Bridges Prokaryotic and Eukaryotic Regulatory Paradigms

**DOI:** 10.1101/2025.05.05.652248

**Authors:** Fernando Medina Ferrer, Dipti D. Nayak

## Abstract

Archaeal transcription is a hybrid of eukaryotic and bacterial features: a RNAP II-like polymerase transcribes genes organized in circular chromosomes within cells devoid of a nucleus. Consequently, archaeal genomes are depleted in canonical transcription factors (TFs) found in other domains of life. Here we outline the discovery of a cryptic archaea-specific family of ligand-binding TFs, called AmzR (<u>A</u>rchaeal <u>M</u>etabolite-sensing <u>Z</u>ipper-like <u>R</u>egulators). We identify AmzR through an evolution-based genetic screen and show that it is a negative regulator of methanogenic growth on methylamines in the model archaeon, *Methanosarcina acetivorans*. AmzR binds its target promoters as an oligomer using paired basic α-helices akin to eukaryotic leucine zippers. AmzR also binds methylamines, which reduces its DNA-binding affinity, and allows it to function as a one-component system commonly found in prokaryotes, while containing a eukaryotic-like DNA-binding motif. The AmzR family of TFs are widespread in archaea and broadens the scope of innovations at the prokaryote-eukaryote interface.

## Introduction

The amalgamation of prokaryotic morphology and eukaryotic information processing machinery in archaea has likely led to the evolution of traits that are unique to this domain of life^1^. Given the abundance and relevance of archaea to humans and the planet, these traits are critical to understand the past, present and future of life on Earth^2,3^. However, a paucity of molecular studies with archaea has impeded the discovery of these features. Distinctly archaeal traits are best exemplified in the context of transcriptional regulation. Archaea use homologs of the eukaryotic TATA-binding protein and transcription factor B to form a preinitiation complex that recruits a RNAP II-like polymerase for transcription initiation^4,5^. However, their genetic material is organized in circular chromosomes and the cell is devoid of a nucleus like bacteria. Thus, transcription regulation in archaea would likely require regulatory proteins that recognize eukaryotic-like promoter elements in the context of a prokaryotic cell. Although several families of regulatory transcription factors (TFs) from bacteria have been identified in archaea, archaeal genomes are still significantly depleted of TFs^6–8^, and there are two likely explanations for this conundrum. One is that genes in archaea are mostly under the control of post-transcriptional and post-translational regulation^9,10^. Alternatively, there are divergent archaea-specific TFs that have evaded discovery through contemporary *in silico* approaches^11^. In support of the latter, here we describe an archaea-specific family of TFs called AmzR that function as a prokaryotic one-component system (OCS) using a eukaryotic DNA-binding motif to regulate methanogenesis.

Methanogenesis—energy conservation coupled to methane production—is a metabolic lifestyle unique to, and widespread within, members of the archaea^12^. Although methanogens play a critical role in anaerobic ecosystems by facilitating the decomposition of organic matter, little is known about how these organisms sense and respond to nutritional cues^13^. We used methanogens within the genus *Methanosarcina* as a model system to explore archaeal gene regulation because: *a)* they are metabolically diverse and their substrate range spans from H_2_+CO_2_ to small organic compounds^14,15^, *b)* they regulate the transcription of enzymes for methanogenesis in response to substrate availability through as yet unknown mechanisms^16–18^, and *c)* they are genetically tractable^19,20^. In *Methanosarcina* spp., substrate-specific methyltransferases (henceforth referred to as MT1s) catalyze the first step of growth on methylated compounds like methanol or methylamines^21–26^. MT1s are among the most highly regulated genes in archaea; their expression can change as much as 1000-fold depending on the presence of the cognate substrate (Fig. S1)^18,27,28^. Prior work suggests that the transcription of certain MT1s is, in part, regulated by Msr proteins, which belong to the ArsR family of regulators with a conserved helix-turn-helix DNA-binding domain^29,30^. Internal or external cues that control Msr activity in response to the cognate substrate are yet to be identified. Similarly, two sensor kinases from *Methanosarcina acetivorans* have been shown to bind methylated sulfur compounds *in vitro*^31,32^, but their regulatory role *in vivo* is yet to be determined. Despite continual efforts to investigate the regulation of methylotrophic methanogenesis in *Methanosarcina* spp., a signal transduction cascade that links the environmental signal (e.g. extracellular concentration of a methylated compound) to a transcriptional response (e.g. expression of MT1s) has not been identified^13^.

In this study, we describe the AmzR family of ligand-binding TFs that regulate methanogenesis in archaea. We use experimental evolution as a forward genetic screen to identify AmzR and its role as a negative regulator of methylotrophic methanogenesis. We provide *in vivo* evidence that AmzR is involved in the transcriptional repression of methylamine-specific MT1s in *M. acetivorans*. *In vitro* analyses reveal that AmzR is a methylamine sensor that also binds its target promoters through paired basic α-helices that resemble eukaryotic DNA-binding motifs like the leucine zipper (bZIP) or the basic helix-loop-helix (bHLH) domains. Altogether, we provide conclusive evidence that AmzR acts as an OCS by tuning the expression of methylamine-specific MT1s in response to methylamine concentrations. Finally, we show that the sequence and function of AmzR is conserved in methanogens that grow on methylamines. Akin to transcription in Archaea—a blend of prokaryotic cell morphology and eukaryotic transcriptional machinery— AmzR functions like a prokaryotic OCS but uses a eukaryotic DNA-binding motif, which could represent an archetype of hybrid gene regulation that is prevalent in this domain of life.

## Results

### Laboratory evolution of *Methanosarcina acetivorans* in an alternating substrate regime targets regulatory genes

Methanogens like *M. acetivorans* can grow on a wide range of methylated compounds by substrate disproportionation, i.e. the methyl group is oxidized to carbon dioxide and reduced to methane in a 3:1 ratio, using a metabolic pathway known as methylotrophic methanogenesis (Fig. 1A). The first step of methylotrophic methanogenesis requires a substrate-specific MT1 whereas the remaining steps are mostly conserved, regardless of the methylated compound used for growth (Fig. 1A). Accordingly, in *M. acetivorans,* we found that the expression of the substrate-specific MT1s for two commonly used methylated compounds—methanol and trimethylamine (TMA)— is substrate-dependent, whereas the rest of the pathway is, by and large, constitutively expressed (Fig. 1A and Fig. S1). Based on this observation, we hypothesized that methanogens would require additional time to rewire MT1 expression when switching from one methylated compound to another, which we refer to as acclimation time henceforth. Indeed, we could reproducibly measure acclimation time in *M. acetivorans* (see Methods) and observed a non-reciprocal trend for methanol and TMA. The acclimation time for the switch from methanol to TMA (*ca.* 96 h) is three-fold longer than from TMA to methanol (*ca.* 36 h) (Fig. 1B). As expected, these acclimation times were consistently greater than the lag times observed during sequential passaging on a given substrate (methanol or TMA), which range from 7 to 18 h (Fig. 1B).

**Figure 1.**
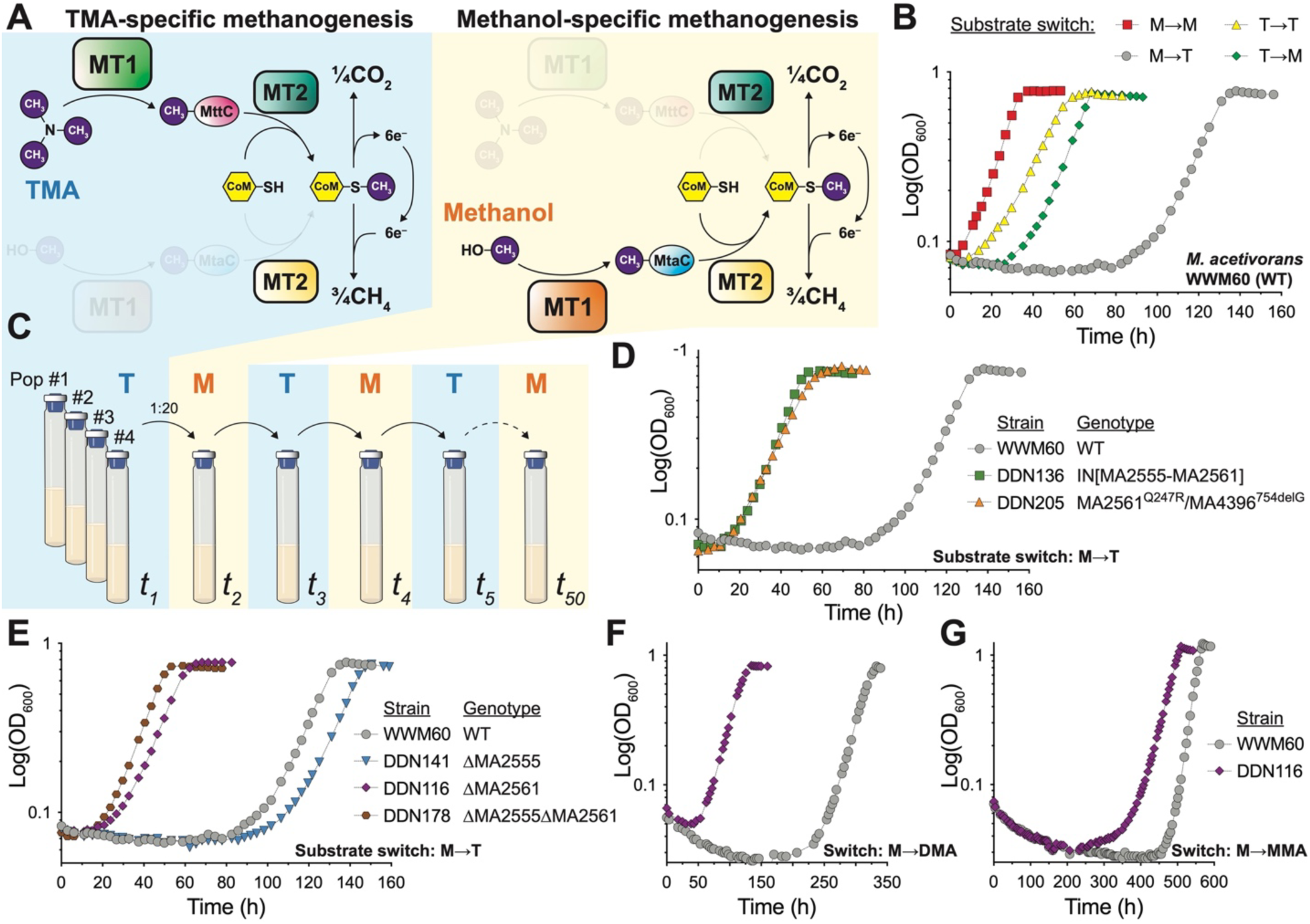
Experimental evolution implicates MA2561 in the regulation of methylamine-mediated methanogenesis in *Methanosarcina acetivorans*. (**A**) During methylotrophic methanogenesis, the expression of a substrate-specific methyltransferase complex (MT1) is tightly regulated by the availability of the corresponding substrate—trimethylamine (TMA; left panel in blue) or methanol (right panel in yellow)—while the expression of other metabolic enzymes remains mostly unchanged (see the entire methylotrophic pathway along with the expression levels of its representative enzymes summarized in Fig. S1). The MT1 complex is comprised of two proteins: a substrate-specific methyltransferase MttB or MtaB, for TMA and methanol respectively, that transfers the methyl group from the substrate (purple sphere) to its cognate corrinoid-binding protein (MttC or MtaC). The methyl group from the corrinoid-binding protein is then transferred to coenzyme M (CoM) by another methyltransferase complex (MT2). The methyl-coenzyme M product generated by MT2 is reduced to methane or oxidized to carbon dioxide in a 3:1 ratio. Additional dimethylamine- and monomethylamine-specific MT1 complexes, not depicted here, are required for the metabolism of dimethylamine (DMA) and monomethylamine (MMA) that are generated during the metabolism of TMA (for more details see Fig. S1). (**B**) Representative growth curves of the *M. acetivorans* parent strain (WWM60) when cells are grown on the same methylated compound as the inoculum (M→M (red) or T→T (yellow) for methanol and TMA, respectively) or after switching to a new methylated compound (M→T (gray) or T→M (green), from methanol to TMA, or from TMA to methanol, respectively). (**C**) A schematic of the evolution experiment conducted with four replicate populations of WWM60 (Pop #1–4) that were alternately grown in minimal media with either TMA (T) or methanol (M) as the sole methanogenesis substrate. During growth on TMA, cells primarily express the methylamine-specific MT1s (left panel in A). Similarly, in media with methanol, cells primarily express methanol-specific MT1s (right panel in A). The evolution experiment comprised of a total of 50 serial transfers (*t_1_*–*t_50_*) (see Fig. S2 for additional details). (**D**) Representative growth curves of the parent strain (WWM60; gray) and isolates from the final transfer (*t_50_*) of Population #1 and Population #4 (DDN136 and DDN205, respectively) when methanol-grown cells were inoculated in medium with TMA (M→T). DDN136 (green) has a genomic inversion that disrupts the coding sequence of MA2555 and MA2561 (IN[MA2555-MA2561]) and DDN205 (orange) has a non-synonymous point mutation in MA2561 and a frameshift mutation in MA4396 (MA2561^Q247R^/MA4396^754delG^). Quantification and statistical analysis of the growth parameters from triplicate growth curves are shown in Fig. S4. (**E**) Representative growth curves of the parent strain (WWM60; gray), ΔMA2555 (DDN141; blue), ΔMA2561 (DDN116; purple), and ΔMA2555ΔM2561 (DDN178; brown) when methanol-grown cells are transferred to TMA-containing media (M→T). Quantification of growth parameters from triplicate growth curves are shown in Figures S7 and S8. Representative growth curves of the parent strain (WWM60; gray) and the ΔMA2561 mutant (DDN116; purple) when methanol-grown cells are transferred to (**F**) dimethylamine (M→DMA) or (**G**) monomethylamine (M→MMA). Quantification and statistical comparison of growth parameters are shown in Figure S9.

We devised an evolution-based forward genetic screen to identify regulatory processes that control the acclimation time during the substrate switch from methanol to TMA and vice versa. Briefly, we initiated four replicate populations of *M. acetivorans* (strain WWM60^33^) in minimal medium with 25 mM TMA as the sole growth substrate. After four days of growth at 37 °C, we transferred a 20-fold dilution of the TMA-outgrowth to minimal medium with 75 mM methanol. After another four days at 37 °C, the methanol-outgrowth was inoculated into fresh medium with 25 mM TMA. This alternating substrate regime was repeated for a total of 50 transfers over the course of 200 days (Fig. 1C and Fig. S2). Since the transition from methanol to TMA (*ca.* 96 h) is the major bottleneck in this evolution experiment, we expected our screen to identify mutants with shorter acclimation times for this switch. Indeed, isolates from the final transfer (*t_50_*) of each evolved population could switch from methanol to TMA in *ca.* 17 h, which is 5.6-fold faster than the parent strain (Fig. 1D, Fig. S3, Fig. S4). Beyond acclimation time, we did not detect any substantial changes in the growth phenotype on methanol, TMA or a combination of methanol and TMA (Fig. S3, Fig. S4). Next, we sequenced the final transfer (*t_50_*) of the four replicate populations to identify underlying mutations. A 4,513-bp genomic inversion disrupting the coding sequence of two genes of unknown function, MA2555 and MA2561, was fixed in three of the four populations. The fourth population had a point mutation in MA2561 (Q247R) and a 1-bp frameshift deletion in a gene encoding a putative hydantoinase/oxoprolinase (MA4396) (Table S1 and Fig. S5). Altogether, our evolution-based genetic screen targeted three distinct loci that might be involved in the regulation of methylotrophic methanogenesis in *M. acetivorans*.

### MA2561 is a negative regulator of methylamine growth in *Methanosarcina acetivorans*

None of the genes targeted in the aforementioned evolution experiment have been studied before, but two of them (MA2555 and MA2561) contain domains commonly found in sensor kinases or other signaling proteins^34,35^ that made us pursue them further (Fig. S6). To evaluate their role in *M. acetivorans* we generated markerless knockouts in the parent strain (WWM60) using our well-established CRISPR technology^19^. We performed whole-genome sequencing of the mutants to verify the absence of suppressors or off-target mutations from CRISPR-editing (Table S2). Like the evolved isolates, the ΔMA2561 and ΔMA2555ΔMA2561 mutants could rapidly switch from methanol to TMA (Fig. 1E; Fig. S7). In contrast, the ΔMA2555 strain took even longer than the parent strain for this substrate switch (Fig. 1E). No other major phenotypic changes were observed for any of these mutants on methanol, TMA, or a combination of both substrates (Fig. S7 and Fig. S8). Based on these data, we conclude that MA2561—the only gene targeted in all four evolved lineages—is responsible for the long acclimation time from methanol to TMA. Since TMA is a composite substrate and produces dimethylamine (DMA) and monomethylamine (MMA) as metabolic intermediates that are consumed by distinct substrate-specific MT1s (Fig. S1), we evaluated the role of MA2561 in regulating acclimation times on these compounds as well. Like TMA, we found that the ΔMA2561 mutant could switch from methanol to DMA or MMA substantially faster than the parent strain (Fig. 1F, Fig 1G, Fig. S9). These data indicate that the regulatory effects of MA2561 span other methylamines too.

To ensure that the phenotype of the ΔMA2561 mutation does not stem from extragenic effects, we complemented the gene *in trans* under a tetracycline-inducible promoter (ΔMA2561::P*mcrB*(tetO4)-MA2561)^33^. Complete induction of MA2561 not only increased the acclimation time for the switch from methanol to TMA but also slowed down growth on TMA substantially (Fig. S10). To evaluate the dose-dependent effect of MA2561 on TMA growth, we tuned its expression by changing the tetracycline concentration in the growth medium. Growth rate and cell yield on TMA were negatively correlated with the expression of MA2561 *in vivo* (Fig. 2A), whereas no such correlation was detectable on methanol or for a control gene (*uidA*) on either substrate (Fig. S11). Overexpression of MA2561 also diminished growth rates on DMA and MMA (Fig. 2B, 2C, and Fig. S12). Multiple attempts to select for suppressor mutations that rescue growth on TMA in the presence of high concentrations of tetracycline ultimately resulted in loss-of-function mutations of MA2561 (Table S3). Altogether, these data provide compelling genetic evidence that MA2561 is a negative regulator of methylamine-mediated methanogenesis *in vivo*.

**Figure 2.**
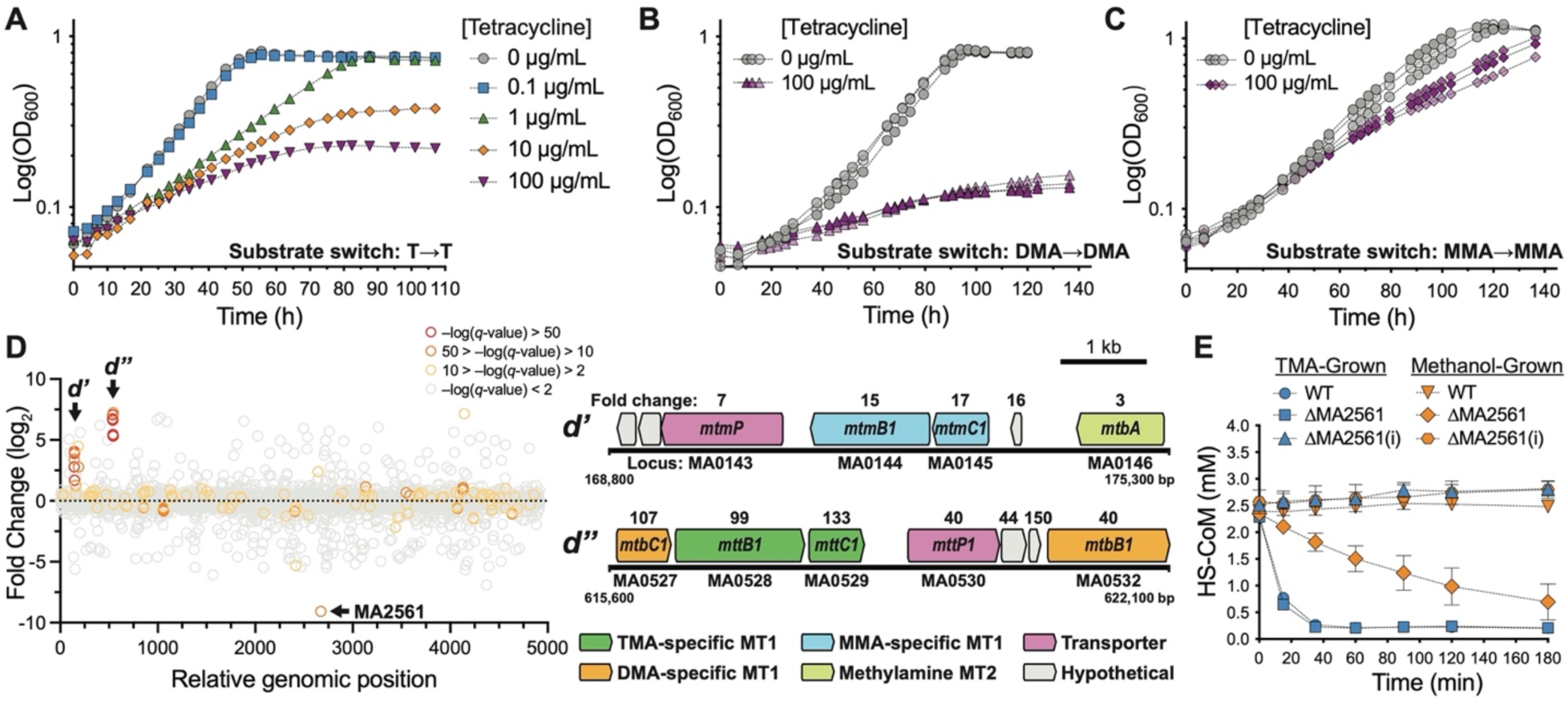
MA2561 is a repressor of methylamine-specific MT1 genes. (**A**) Dose-dependent effect of MA2561 on trimethylamine (TMA) growth. In a mutant strain expressing a single-copy of MA2561 under the control of a tetracycline-inducible promoter [P*mcrB*(tetO4)] (DDN152), its expression can be tuned by changing the concentration of tetracycline in the growth medium (0– 100 µg/mL). Increase in tetracycline concentration impairs growth on TMA. Quantification and statistical comparison of growth parameters are shown in Figures S10 and S11. Overexpression of MA2561 in minimal media containing either (**B**) dimethylamine (DMA), or (**C**) monomethylamine (MMA) also impairs growth. Quantification and statistical comparison of growth parameters are shown in Figure S12. (**D**) Transcriptional profile of the ΔMA2561 mutant compared to its parental strain (WWM60) during growth on methanol ordered by genes based on their location on the chromosome. Genes with a statistically significant [–log_10_(*q*-value) > 2] change in expression are shown by colored circles while genes with non-significant changes are shown in gray. Note that there are two chromosomal loci where most differentially expressed genes are present as indicated by black arrows: *d’* contains the MMA-specific MT1 isoform 1 operon (*mtmCB1*) and *d’’* contains the TMA- and DMA-specific MT1 isoform 1 operon (*mtbCB1– mttCB1*). Fold change values are indicated above each gene in *d’* and *d’’*. (**E**) TMA-specific methyltransferase activity of crude cell lysates measured by the rate of consumption of coenzyme M (HS-CoM) over time in the presence of 100 mM TMA as the reaction substrate. Measurements for cells extracts and their heat-inactivated controls (i) from WWM60 or the ΔMA2561 mutant grown on either TMA (blue) or methanol (orange) are shown. Error bars represent SEM of triplicate assays. Quantification and statistical comparison of the specific activity for each condition is shown in Figure S15.

### MA2561 represses a methylamine-specific metabolic regulon in *Methanosarcina acetivorans*

To understand how MA2561 regulates methylamine growth, we compared the global transcriptome of the ΔMA2561 mutant to the parent strain (WWM60) on methanol and observed that many of the methylamine-specific MT1 genes were highly expressed (by up to 100-fold) in the ΔMA2561 mutant (Fig. 2D, Figs. S13, S14, and Table S4). To test if an increase in the expression of methylamine-specific MT1s corresponds to a concomitant increase in enzyme activity, we performed anaerobic methyltransferase assays with cell lysates. As expected, methanol-grown cell extracts from the ΔMA2561 mutant had higher TMA-methyltransferase activity compared to WWM60 (Fig. 2E), although not to the extent observed during TMA growth (Fig. S15). Accordingly, the expression of methylamine-specific MT1s in the ΔMA2561 mutant was higher on TMA too (Fig. S15). Thus, even though additional modes of post-transcriptional^9,25^ and post-translational^36^ regulation likely exist, our data clearly show that MA2561 is involved in the transcriptional repression of methylamine-specific MT1 genes in *M. acetivorans*.

### MA2561 is an intracellular sensor of methylamines

MA2561 is predicted to be a cytosolic protein with two domains (PAS and GAF) that are implicated in ligand-binding and routinely found in input modules of signaling proteins^34,35^ (Fig. S6). Therefore, we hypothesized that MA2561 might regulate the expression of methylamine-specific MT1s by acting as an intracellular sensor of methylamines. To test this hypothesis, we introduced an affinity purification tag at the C-terminus of the MA2561 CDS and expressed it under the control of a tetracycline-inducible promoter in the ΔMA2561 background (Fig. S16). Upon induction, the tagged protein was detectable by Western Blot and inhibited growth on TMA (Fig. S16), which indicates that the sequence and location of the tag does not impact the function of MA2561 *in vivo*. Affinity purification of tagged MA2561 results in a soluble product that produces a single ∼50 kDa band on a reducing SDS-PAGE gel (Fig. S17). Using differential scanning fluorimetry, we observed that the thermal stability of the purified protein increased substantially in the presence of methylamines (Fig. 3A and S18). This thermal shift reflects binding of methylamines to MA2561, which was not detected in the presence of methanol, or for a control protein in the presence of methylamines (Fig. S18). Overall, we have shown that MA2561 operates as a methylamine sensor for the regulation of the cognate MT1 genes.

**Figure 3.**
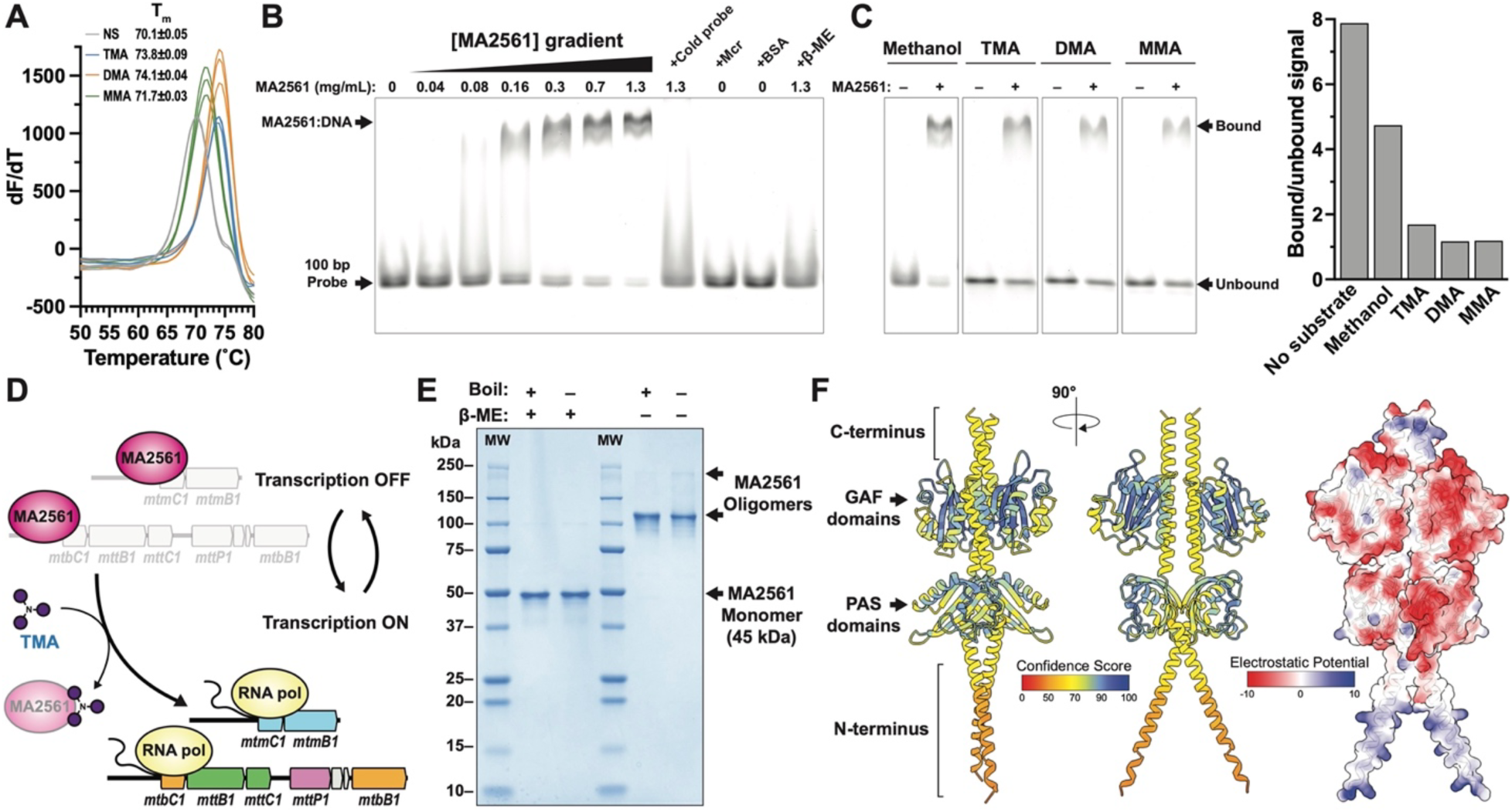
MA2561 binds methylamines and the promoter region of monomethylamine-specific MT1. **(A)** Differential scanning fluorimetry of purified MA2561 shows a ∼2–4 °C thermal shift upon incubation with trimethylamine (TMA; blue), dimethylamine (DMA; orange), and monomethylamine (MMA; green) relative to MA2561 without any substrate (NS; gray). Derivatives of triplicate melt curves and their T_m_ are shown for each condition. No significant thermal shift was observed with methanol, or for a control protein in the presence of methylamines (see Figure S18). (**B**) Electrophoretic mobility shift assay (EMSA) using MA2561 and a 100-bp Cy5-labeled probe containing the promoter elements of the MMA-specific MT1 upregulated in the ΔMA2561 mutant (*mtmCB1* operon) (see Figure S22 for additional details). The formation of a MA2561:DNA complex, as indicated by a gel shift of the 100 bp DNA probe, intensifies with an increase in [MA2561]. This gel shift does not occur in the presence of an excess amount of unlabeled probe (“cold probe”) or if MA2561 is treated with the reducing agent β-mercaptoethanol (β-ME). No gel shift is observed for the DNA probe with other proteins, such as methyl-coenzyme M reductase (Mcr) or bovine serum albumin (BSA). EMSA assays with MA2561 for additional probes spanning the *mtmCB1* promoter region or using other *M. acetivorans* proteins are shown in Figure S22. (**C**) EMSA of the 100-bp probe from **B** with 1.3 mg/mL of MA2561 in the presence of trimethylamine (TMA), dimethylamine (DMA), monomethylamine (MMA), and methanol. Quantification of the ratio of bound and unbound probe (MA2561^+^ lane) shows a decrease of the DNA:protein complex in the presence of methylamines (right graph). (**D**) Proposed model for regulation by MA2561. In the absence of methylamines, MA2561 tightly binds to and occludes the promoter of *mtmCB1*, *mtbCB1*, and *mttCB1* genes, thus preventing their expression. When methylamines, like TMA are present, they bind to MA2561, which leads to conformational changes that ultimately reduce the affinity of MA2561 for its target DNA relative to the transcription initiation complex. Consequently, the promoters of the *mtmCB1*, *mtbCB1*, and *mttCB1* genes are more available for transcription by the RNA polymerase. (**E**) SDS-PAGE of MA2561 under non-reducing conditions (without β-ME) shows the appearance of a primary band at ∼116 kDa and a secondary band at ∼235 kDa that likely correspond to the dimer and the tetramer, respectively (see more details in Figure S23). (**F**) AlphaFold predictions of a MA2561 dimer shown as a ribbon diagram (left and center panel) and a space-filling model colored by surface electrostatic potential (right panel). Positively charged α-helices at the N-terminus form a basic and flexible coiled-coil domain that could potentially bind DNA, analogous to the eukaryotic basic leucine zipper (bZIP) and basic helix-loop-helix (bHLH) DNA-binding domains as shown in Figure S24.

### MA2561 is a one-component system that controls the expression of methylamine-specific MT1s in response to methylamine concentrations

Even though MA2561 binds methylamines, no obvious DNA-binding motifs are present in its primary sequence (Fig. S6). Therefore, we initially hypothesized that MA2561 might participate in a signal transduction cascade that involves at least one other DNA-binding response regulator. To the contrary, we found overwhelming evidence that MA2561 is singularly involved in the regulation of methylamine-specific MT1s. First, we were unable to identify any potential interaction partners in pull-down assays with MA2561 using lysates from either methanol or TMA grown cells (Fig. S17). Next, we tested if MA2561 might participate in regulation via phosphorelay akin to a histidine kinase and response regulator in a two-component system^10,37^ by measuring its phosphorylation state *in vivo.* We could not detect any phosphorylated protein during growth on methanol or methylamines (Fig. S19). We also conducted a second round of substrate-switching evolution experiments, reasoning that mutations in genes other than MA2561 might help us identify additional proteins that participate in the same signal transduction cascade (Fig. 1C). However, all ten independently derived evolved populations contained loss-of-function mutations in MA2561 (Table S5). Finally, we tested the potential role of MsrB in this signaling pathway, as prior work has reported that a Δ*msrB* mutant constitutively expresses methylamine-specific MT1s^38^. To this end, we generated a Δ*msrB* mutant in the WWM60 genetic background (Table S2) and found that it took even longer than WWM60 to switch from methanol to TMA (Fig. S20). We also conducted an epistasis test by expressing MA2561 under the control of the tetracycline-inducible promoter in the Δ*msrB* mutant. If MA2561 represses the methylamine-specific MT1s through MsrB, the absence of MsrB would preclude the detrimental effect of MA2561 overexpression during TMA growth (Fig. 2A). In contrast, we still observed a dose-dependent, TMA-specific growth inhibition by MA2561, which indicates that MsrB and MA2561 mediated-regulation do not crosstalk *in vivo* (Fig. S21). Taken together, all these independently derived lines of evidence suggest that MA2561 might not participate in a multi-component signal transduction cascade *in vivo.* Thus, we entertained the possibility that MA2561 could directly repress the target MT1s, by forming a cryptic DNA-binding motif that cannot be detected *in silico*.

To assess the DNA-binding properties of MA2561, we conducted gel-shift assays with the promoter and 5’-untranslated regions (UTR) of the MMA-specific MT1 (*mtmCB1*) that is upregulated in the ΔMA2561 mutant. Our choice of the *mtmCB1* promoter was motivated by conclusive evidence of its location based on *in vitro* studies unlike any other methylamine-specific MT1 promoter in *Methanosarcina* spp.^39^ (Fig. S22). By using DNA probes of varying lengths, we narrowed in on a 100-bp region containing the TATA box, the potential B-recognition element (BRE), and the transcriptional start site (TSS) of *mtmCB1* that is, both, necessary and sufficient for MA2561 binding (Fig. 3B and Fig. S22). This region does not bind bovine serum albumin (BSA) or other proteins from *M. acetivorans*, and a fragment upstream of this region does not bind MA2561, suggesting that the interaction is sequence- and protein-specific (Fig. S22). We also observed that the addition of methylamines, but not methanol, attenuates complex formation between MA2561 and the *mtmCB1* promoter (Fig. 3C). Altogether, these data are consistent with MA2561 acting as an OCS that binds DNA and occludes the promoter of methylamine-specific MT1 genes in the absence of methylamines (Fig. 3D).

### Oligomeric MA2561 forms a basic coiled-coil DNA-binding motif

Signal transduction by OCS is widely distributed in Bacteria and Archaea^40,41^, including among others the DtxR^42^, TetR^43^, TmrB^44,45^, LysR^46^, Lrp^47,48^, Lrs14^49,50^, MarR^51^, NrpR^52^, and SmtB/ArsR^53^ TF families, all of which contain a helix-turn-helix (HTH) DNA-binding domain unlike MA2561. While the MA2561 monomer lacks a canonical DNA-binding domain (Fig. S6), we hypothesized that a higher-order oligomer could, perhaps, present a cryptic DNA-binding motif. To test this hypothesis, we visualized the purified protein under non-reducing conditions and noted the appearance of high molecular weight bands, with the most dominant species appearing at *ca.* 116 kDa that likely corresponds to a dimer (Fig. 3E, Fig. S23 and Table S6). We used Ellman’s reagent to quantify that at least one cysteine residue per monomer forms a disulfide bridge in anaerobic preparations of MA2561 (Fig. S23). Given the proximity and orientation of Cys^3^^24^ in Alphafold predictions^54^, it is likely that this residue is vital for interchain disulfide bridges that lead to the formation of a MA2561 dimer *in vivo.* To test if dimerization is crucial for DNA-binding, we conducted gel-shift assays with fully reduced MA2561 and found that it could no longer bind the 100-bp region containing the promoter elements of *mtmCB1* (Fig. 3B). Knowing that oligomeric MA2561 is critical for DNA biding, we examined the electrostatic surface potential in structural models of dimeric MA2561 to locate the putative DNA-binding region. While the surface of the PAS and GAF domains are predominantly acidic, and thus unlikely to engage in direct contacts with DNA, both the N- and C-termini contain positively charged α-helices that are packed in a parallel arrangement resembling a coiled-coil domain (Fig. 3F). The C-terminal helices contain a cluster of basic amino acids at the tips, but this motif appears too rigid and small to accommodate DNA without a significant structural rearrangement of its adjacent GAF domains. In contrast, the N-terminal helices form a flexible coiled-coil-like domain containing several basic residues that could grip DNA in a fashion reminiscent to basic leucine zipper (bZIP)^55^ and basic helix-loop-helix (bHLH)^56^ DNA-binding motifs found in eukaryotic TFs (Fig. S24). Consistent with its putative role in DNA-binding, many basic residues in the N-terminus are conserved across MA2561 homologs (Fig. S24). Altogether, we provide strong evidence of MA2561 dimerization into a hitherto cryptic DNA-binding domain formed by positively charged α-helices at the N-terminus. The striking structural resemblance of this domain to eukaryotic bZIP and bHLH motifs is in stark contrast to the helix-turn-helix (HTH) DNA-binding domain architecture found in most other prokaryotic OCSs^7,40,41,57^.

### Functional AmzR family regulators are prevalent in archaea

Since methylamine metabolism is broadly distributed in methanogens, we wanted to evaluate whether the underlying regulatory processes might be conserved as well. To this end, we looked at the distribution of MA2561 homologs in methanogens within the *Methanosarcinia*, a large group of methanogens that contains nearly all cultured isolates capable of methylamine growth. Indeed, the presence of MA2561 homologs in *Methanosarcinia* strongly correlates with the presence of methylamine-specific MT1 genes (P<0.0001, Fisher’s exact test; Fig. 4A and Table S7). Most methylamine-metabolizing methanogens encode at least one copy of a MA2561 homolog that is likely vertically inherited (Fig. 4A and Fig. S25). Notably, the correlation between MA2561 and methylamine metabolism is especially well-conserved in free-living methanogens. In contrast, host-associated methanogens, like *Methanimicrococcus* and *Methanolapillus* spp. which live in the gut of arthropods, seem to have lost MA2561-type regulators^58^ (Fig. 4A). This observation is consistent with genome streamlining accompanied by a loss of TFs in microorganisms that are host-associated or have a parasitic lifestyle^7,59–61^.

**Figure 4.**
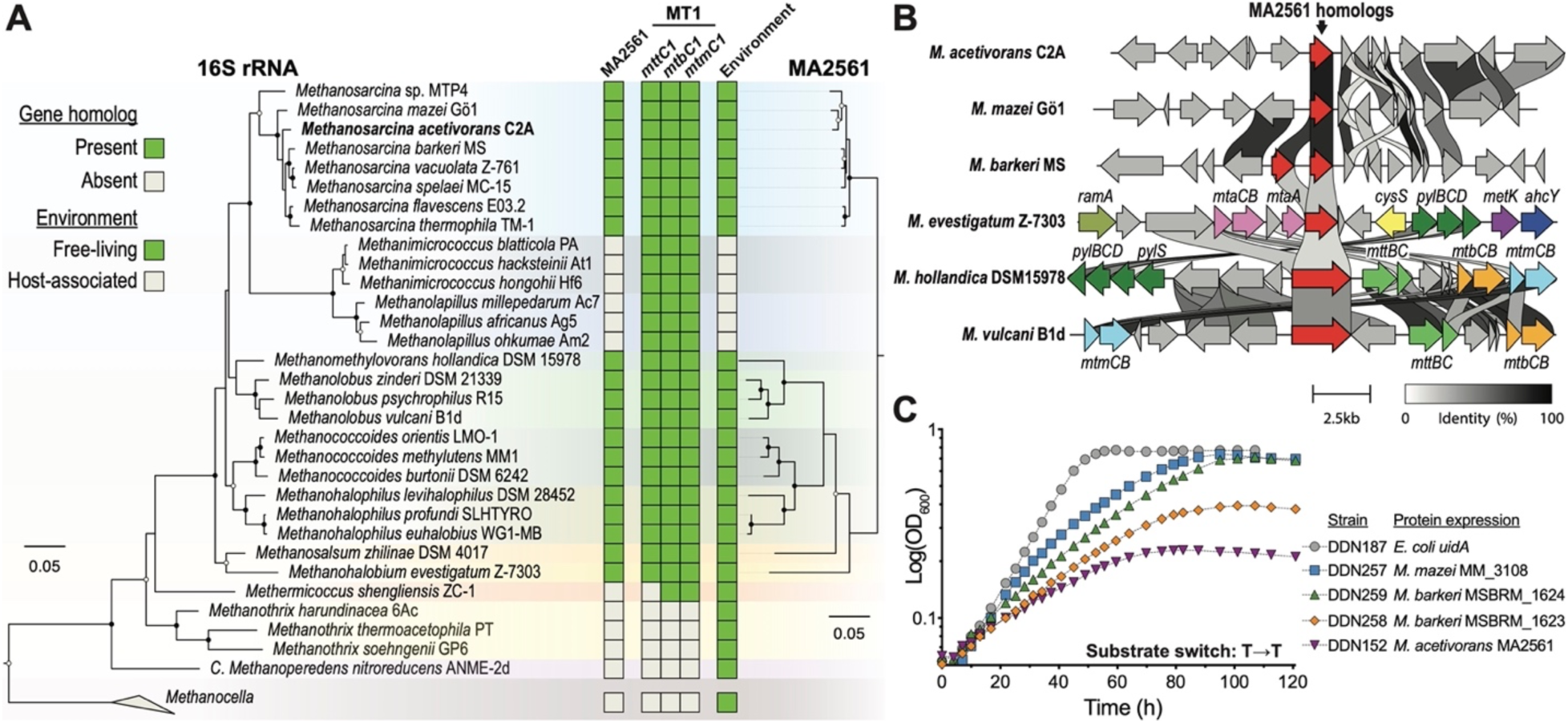
MA2561 homologs are widely conserved in free-living methylamine metabolizing methanogens. (**A**) Maximum likelihood tree of the 16S rRNA locus (left) and MA2561 homologs (right) in members of the Class *Methanosarcinia*. The distribution of MA2561 homologs and trimethylamine-, dimethylamine- and monomethylamine-specific MT1 genes (*mttC*, *mtbC*, and *mtmC,* respectively) are shown (green: present; gray: absent). In addition, the environmental distribution of the strains is also shown (green: free-living; gray: host-associated). For the phylogenetic trees, bootstrap support >70 is shown at the nodes (70–90, gray circles; >90, black circles). (**B**) Genomic context of MA2561 and its homologs (in red) in select strains. MA2561 homologs are often found in the vicinity of important genes involved in methylotrophic methanogenesis in members of the *Methanohalobium*, *Methanomethylovorans*, and *Methanolobus* spp. Genes of unknown function are shown in gray. Known functional annotations are as follows: *mtaCB* (pink): methanol-specific MT1, comprised of the corrinoid-binding protein *mtaC* and the methanol:5-hydroxy-benzimidazolyl-cobamide methyltransferase *mtaB*; *mtaA* (pink): methylcobamide:coenzyme M methyltransferase MT2; *mttCB* (light blue): trimethylamine (TMA)-specific MT1; *mtbCB* (orange): dimethylamine (DMA-specific MT1; *mtmCB* (light green): monomethylamine (MMA)-specific MT1; *ramA* (olive): reductive activation of methyltransfer, amines; *pylBCD* (dark green): pyrrolysine biosynthetic machinery; *pylS*: pyrrolysine—tRNA ligase; *cysS* (yellow): cysteine—tRNA ligase; *metK* (purple): methionine adenosyltransferase; *ahcY* (blue): adenosylhomocysteinase. (**C**) Representative growth curves of the ΔMA2561 mutant with a vector expressing MA2561 homologs from *M. mazei* Gö1 (MM_3108; blue) or *M. barkeri* MS (MSBRM_1623 and MSBRM_1624; orange and green respectively) under the control of the tetracycline inducible promoter P*mcrB*(tetO4). All strains were grown in minimal media supplemented with 25 mM TMA and 100 µg/ml tetracycline. Growth of cells expressing either *uidA* (gray) or MA2561 (purple) are shown as negative and positive controls, respectively. Quantification and statistical analysis of growth parameters, along with control growth curves in the absence of tetracycline or on methanol are summarized in Figure S26.

Even though MA2561 is not located in an operon or near the genes it regulates in *M. acetivorans*, it is often found near methylamine-specific MT1s, putative methylamine transporters, and the *pylBCD* operon (pyrrolysine biosynthetic machinery) in *Methanohalobium, Methanomethylovorans,* and *Methanolobus* spp. (Fig. 4B). The proximity of MA2561 homologs to genes that are critical for methylamine metabolism further reinforces our proposed function. To test if the role we identified for MA2561 in *M. acetivorans* is also conserved in other methanogens, we expressed a homolog from *M. mazei* Gö1 (MM_3108), and two from *M. barkeri* MS (MSBRM_1623 and MSBRM_1624) under the control of the tetracycline-inducible promoter in the ΔMA2561 mutant. As observed for MA2561 (Fig. 2A), all these homologs lead to a dose-dependent growth inhibition on TMA (Fig. 4C), but not on methanol (Figure S26). Notably, both paralogs of *M. barkeri* repress methanogenic growth, albeit with different relative strengths (Fig. S26). The structural models of these homologs also contain a putative DNA-binding region comprised of paired basic α-helices at the N-terminal region (Fig. S27). These data provide evolutionary support for the *in vivo* role of MA2561 as an OCS with a widely conserved function across methylamine-metabolizing methanogens. Altogether, based on the function that we established for MA2561 and its homologs, we have named this locus *amzR* (<u>A</u>rchaeal <u>M</u>etabolite-sensing <u>Z</u>ipper-like <u>R</u>egulator).

## Discussion

This study outlines the discovery of an archaeal family of OCSs—AmzR and its homologs—that regulate methanogenesis through the formation of a basic DNA-binding motif akin to eukaryotic bZIP and bHLH TFs. While our analysis specifically dives into the role of AmzR in the context of methylamine metabolism, we expect that the AmzR-type family of regulators has diverse regulatory roles across methanogens and other archaea. *M. acetivorans* alone codes for three additional PAS_n_-GAF AmzR-like proteins of unknown function and multiple other regulators with PAS and GAF domains but no discernable output domains (Table S8). Moreover, these PAS and GAF domain containing proteins are enriched across archaeal genomes too^10,37,62^. Since the ability of AmzR to bind DNA relies on its oligomeric state (Fig 3B), it is possible that the formation of both homo and heterodimeric species allow the cell to produce a combinatorial bonanza of proteins with varying DNA-binding specificities, broadening the regulatory landscape of AmzR-type regulators. Indeed, a similar mechanism is well documented in eukaryotes for bZIP and bHLH TFs^63–65^, where oligomerization, to form homo and heterodimers, leads to complex regulatory networks that allow tight control over diverse processes, from development and cellular differentiation^66,67^ to neurodegenerative diseases^68^ and cancer^69^.

For decades, genome-wide surveys have noted a conspicuous deficit of TFs in archaeal genomes^6,8,11^ that are currently estimated to be at par with the low density of TFs observed in parasitic and pathogenic bacteria^7^. Contrary to bacteria, the number of archaeal TFs does not necessarily scale with genome size or metabolic diversity either, especially for *Methanosarcina* spp. that have some of the largest archaeal genomes and are among the most metabolically versatile methanogens^6,11,70^. In this work we show that the AmzR family of OCSs account, in part, for some of the “missing” archaeal TFs. Furthermore, both archaeal and eukaryotic genomes are depleted in two-component systems (TCS) compared to bacteria^6,71^. While the shortage of TCS in eukaryotes is balanced by other systems such as G protein-coupled receptors, the scarcity—or in some cases complete absence—of known TCS in archaea is puzzling^10,71^, especially considering that they often encode response regulators that lack canonical output domains^40,62,72^. Many of these radically different response regulators of unknown function contain PAS_n_-GAF domain architectures that resemble AmzR^10,37,62^ (Table S8). Thus, the cryptic AmzR-type DNA-binding motif may also account for the missing output domains in archaeal TCS. Beyond methanogenesis, we anticipate that the AmzR family of TFs regulate a multitude of features across archaea using DNA-binding motifs analogous to bZIP and bHLH, possibly representing living fossils of early eukaryotic transcriptional regulation.

## Conclusion

Mechanisms involved in the regulation of methanogenesis have long been elusive. Here, in our quest to identify transcriptional regulators of methanogenesis, we uncovered AmzR, a uniquely archaeal innovation at the prokaryote-eukaryotic interface. On the one hand, AmzR exemplifies a common prokaryotic strategy to control transcription using OCSs. On the other, AmzR has a DNA-binding domain verisimilar to the eukaryotic bZIP and bHLH motifs. Neither the form or the function of this protein could be predicted from its primary sequence, and through rigorous genetic and biochemical assessment we have identified a previously unaccounted family of TFs in Archaea.

## Materials and Methods

### Culture conditions

*M. acetivorans* strains were cultivated at 37 °C in a Heratherm oven (Thermo Fisher Scientific, Waltham, MA, USA), without shaking in bicarbonate-buffered high salt (HS) medium with N_2_/CO_2_ (80:20) at 55-70 kPa in the headspace as described previously^73^. For protein purification experiments, cells were grown in 500 mL HS medium in 1 L anaerobic bottles sealed with butyl rubber stoppers (Chemglass Life Sciences, Vineland, NJ, USA) containing either 150 mM methanol or 50 mM trimethylamine (TMA) as the substrate along with 2 µg/mL puromycin (to maintain the expression vector encoding the target gene) and 100 µg/mL tetracycline (to induce gene expression)^33^. For growth curves, transcriptomics, and evolution experiments, cells were grown in 10 mL HS medium in Balch tubes sealed with butyl rubber stoppers containing either 75 mM methanol, 25 mM TMA, 50 mM dimethylamine (DMA), 150 mM monomethylamine (MMA) or a combination of 12.5 mM TMA and 37.5 mM methanol. Aliquots of anaerobic puromycin to a final concentration of 2 µg/mL and of fresh anaerobic tetracycline hydrochloride solution to the indicated concentration were added for growth experiments of *M. acetivorans* strains containing expression vectors.

### Laboratory Evolution Experiments

For the first round of laboratory evolution experiments, four replicate populations of the *M. acetivorans* strain WWM60^33^ were cultivated in HS media with 25 mM TMA at 37 °C for four days, after which a 1:20 dilution of the outgrowth was transferred to fresh HS media with 75 mM methanol and incubated at 37 °C for four days. A 1:20 dilution of the methanol outgrowth was transferred to fresh HS media with 25 mM TMA. This substrate-switching regime was continued for a total of 50 transfers. For the second round of evolution experiments, 10 single-colony isolates of WWM60 were passaged in the same substrate-switching regime described above except that cells were transferred every three days for a total of 20 transfers.

### Growth assays and growth parameter calculations

All growth experiments with *M. acetivorans* were conducted by inoculating a 20-fold dilution of a late-exponential culture in 10 mL HS medium with the indicated growth substrate. All inoculations were conducted in an anaerobic atmosphere (*ca*. 78:18:4 N_2_:CO_2_:H_2_, Coy Lab Products, Grass Lake, MI, USA) using a 1 mL syringe. Growth was followed by measuring optical density readings at 600 nm (OD_600_) using a UV-Vis spectrophotometer (GENESYS 50, Thermo Fisher Scientific, Waltham, MA, USA) outfitted with an adjustable single test tube holder. Growth curves were visualized on a semi-logarithmic plot and specific growth rates were obtained from the slope of a linear regression (> 0.99 R^2^ values) using at least 7 timepoints in the exponential phase of growth. Lag and acclimation times were defined as the time (h) at which the linear regression for exponential growth (described above) intersects the y=ln(initial OD_600_) line in the semi-logarithmic plot of each individual growth curve. Growth yield was defined as the maximum OD_600_ value. When cultures reached an OD_600_ reading > 0.95, cells were diluted in HS medium (1:11 dilution) for accurate measurement. All growth experiments were conducted in triplicate and statistical analyses of growth parameters were performed in GraphPad Prism v10.3.1 using unpaired *t*-tests with Welch’s correction.

### Plasmid construction and generation of CRISPR-edited mutants

Twenty-base pair (bp) long sgRNA sequences targeting the MA2555 (MA_RS13285), MA2561 (MA_RS13315), or *msrB* (MA0460; MA_RS02405) CDS were designed *in silico* using the CRISPR site finder tool in Geneious Prime v.11.0 using an NGG-3’ PAM site and including the *M. acetivorans* C2A genome and the pDN201 sequence to screen for off-target matches. The 20-bp sgRNA sequences were synthesized as 5’ overhangs on primers (Integrated DNA Technologies, Coralville, IA, USA) used for introducing the sgRNA expression cassette in pDN201 linearized with *AscI* using Gibson assembly as described previously^19,74^. A ∼2 kilobase pair (kbp) homology repair template sequence (∼1 kbp upstream and ∼1 kbp downstream) to generate an in-frame deletion of the target gene was introduced into the vector with the corresponding sgRNA using Gibson assembly as described previously^19^. Plasmids used for complementation and controlled gene expression under the tetracycline-inducible promoter P*mcrB(tetO4)* were constructed by Gibson assembly using the pJK029A plasmid as backbone^33^. All pDN201- and pJK029A-derived plasmids were retrofitted with the pAMG40^33^ using the Invitrogen Gateway BP Clonase II Enzyme mix (Thermo Fisher Scientific, Waltham, MA, USA) to generate the final cointegrate that can replicate autonomously in *Methanosarcina* as described previously^33^.

All plasmids were introduced in *E. coli* WM4489 as the host strain by electroporation (MicroPulser Electroporator, Bio-Rad, Hercules, CA, USA). All pDN201- and pJK029A-based plasmid containing strains were plated on lysogeny broth (LB)-agar at 37 °C supplemented with 10 µg/mL chloramphenicol. All pAMG40 cointegrates were plated on LB-agar at 37 °C supplemented with 10 µg/mL chloramphenicol and 50 µg/mL kanamycin. For plasmid extraction, cells were grown shaking at 250 rpm in an incubator (Thermo Fisher Scientific, Waltham, MA, USA) at 37 °C in test tubes with 5 mL LB medium supplemented with the corresponding antibiotic(s) and 10 mM rhamnose to increase the plasmid copy number as described before^75^. Plasmid extractions were conducted using the Zymo Zyppy Plasmid Miniprep kit (Zymo Research, Irvine, CA, USA). The pDN201- and pJK029A-derived plasmid sequence was verified by PCR and Sanger sequencing at the UC Berkeley DNA Sequencing Facility. All pAMG40 cointegrates were verified by endonuclease restriction assays.

Transformation of *M. acetivorans* was conducted with 20 mL cultures of the corresponding strain at mid-exponential phase grown in HS medium with 50 mM TMA. Cells were pelleted and lipofected inside an anaerobic chamber (*ca*. 78:18:4 N_2_:CO_2_:H_2_ composition, Coy Lab Products, Grass Lake, MI, USA) with 2 µg of plasmid as described previously. Cells were plated on agar-solidified HS medium containing 50 mM TMA and 2 µg/mL puromycin and incubated in an intra-chamber Wolfe incubator^77^ at ∼37 °C under a 79.9:20:0.1 N_2_:CO_2_:H_2_S atmosphere as previously described^20^. Puromycin-resistant transformants were screened for chromosomal mutations by PCR (for CRISPR editing) or a gene on the expression vector (for pJK029A-derived plasmids). Colonies that tested positive for chromosomal mutations by CRISPR editing were struck on HS media with 50 mM TMA and 20 µg/mL of the purine analog 8-aza-2,6-diaminopurine (8-ADP; CarboSynth, San Diego, CA) for counterselection of the CRISPR-editing plasmid as described previously^78^. Colonies that tested positive pJK029A-derived plasmids were grown in liquid HS medium supplemented with 50 mM TMA and 2 µg/mL puromycin. CRISPR-edited mutants were grown in 10 mL HS medium with 50 mM TMA and 20 µg/mL 8-ADP until the editing plasmid was cured as verified by PCR of the puromycin N-acetyltransferase gene. All primers, plasmids and strains generated and used for this study are listed in Supplementary Tables S9-S11 respectively.

### Genomic DNA extraction and whole-genome sequencing

Genomic DNA was extracted from 6 mL saturated culture of a strain or an evolved population using the QIAGEN DNeasy Blood & Tissue kit (QIAGEN, Hilden, Germany) following the manufacturer’s instructions. Preparation of Illumina DNA library and sequencing was conducted at SeqCenter (Pittsburgh, PA, USA). Sequencing data were analyzed using *breseq* v0.35.5^79^ using population-mode. All raw DNA-sequencing reads have been deposited into the Sequencing Reads Archive (SRA) and the accession number will be made available upon publication.

### RNA extraction and transcriptome analysis

One mL of triplicate cultures in mid-exponential phase (0.3–0.5 OD_600_) were added to 1 mL of pre-warmed (37 °C) TRIzol (Life Technologies, Carlsbad, CA, USA) and incubated at room temperature for 5 min. An equivalent volume of cold (–20 °C) ethanol was then added, and RNA was extracted from 700 µL of the mixture using the QIAGEN RNeasy Mini Kit (QIAGEN, Hilden, Germany) following the manufacturer’s instructions. RNA quality and concentration was assessed using a Nanodrop One UV-Vis Spectrophotometer (Thermo Fisher Scientific, Waltham, MA, USA) before flash freezing at –80 °C. Samples were submitted to SeqCenter (Pittsburgh, PA, USA) for DNAse treatment and library preparation with the Illumina Stranded Total RNA Prep, Ligation with Ribo-Zero Plus kit, using the 10bp IDT for Illumina indexes. An Illumina NextSeq 2000 was used for sequencing paired end 50 bp reads. Demultiplexing, quality control, and adapter trimming was done with BCL Convert v3.9.3. Non-interleaved paired-ended fastq files were uploaded and analyzed in the KBase platform^80^ as previously described^81^. Read quality was evaluated with FastQC v0.11.0 and sequences were aligned with HISAT2 v2.1.0^82^ using the reference genome of *M. acetivorans* C2A^70^. Reads were then assembled with Cufflinks v2.2.1^83^, where the expression value of each gene was obtained as the log_2_(FPKM, fragments per kilobase per million mapped reads). Differential expression values were calculated with DESeq2 v1.20.0^84^ using default parameters, and a *q* < 0.01 was used as the threshold for significant differences. All raw reads from transcriptomic studies conducted for this work have been deposited into the Sequencing Reads Archive (SRA) and the accession number will be made available upon publication.

### Methyltransferase activity assay

Methylamine methyltransferase activity was measured under anaerobic conditions using cell lysates as previously described^24,25,85^ with modifications. Cells were harvested from 20 mL mid-exponential phase cultures in HS media with either 50 mM of TMA or 150 mM methanol by centrifugation at 1,500 ×*g* for 15 min in a E8 Centrifuge (LW Scientific, Lawrenceville, GA, USA) inside an anaerobic chamber with a 96:4 N_2_:H_2_ atmosphere. Cell pellets were washed by gentle resuspension in 10 mL of anaerobic HS-MOPS pH 7.0 (50 mM MOPS pH 7.0, 400 mM NaCl, 13 mM KCl, 54 mM MgCl_2_·6H_2_O, 2 mM CaCl_2_·2H_2_O). Cells were centrifuged again and the pellet was resuspended in 300 µL of hypotonic lysis solution containing 5 mM Ti(III) citrate solution pH 7.0 (added from a 50 mM titanium (III), 86 mM sodium citrate pH 7.0 stock solution prepared as described previously^86^) with 2 U/mL DNase I (Thermo Fisher Scientific, Waltham, MA, USA) and 1X Roche cOmplete EDTA-free Protease Inhibitor Cocktail (MilliporeSigma, Burlington, MA, USA). Lysates were incubated on ice for 1 hour and sonicated for 5 mins in an ultrasonic bath inside the anaerobic chamber. Cell debris were removed by centrifugation at 2,000 ×*g* for 10 min using a Corning LSE mini microcentrifuge inside the anaerobic chamber and the final protein concentration of the cell extracts was quantified using the Pierce Bradford Plus Protein Assay Reagent^87^ (Thermo Fisher Scientific, Waltham, MA, USA). Methyltransferase reactions were carried out inside an anaerobic chamber with a 96:4 N_2_:H_2_ atmosphere composition in 1 mL of 2 mg protein extract, 25 mM MOPS pH 7.0, 2 mM 2-mercaptoethanesulphonic acid (coenzyme M), 3.2 mM 2-bromoethanesulphonic acid, 20 mM MgCl_2_, 0.1 mM ZnSO_4_, 10 mM ATP, and 1.5 mM Ti(III) citrate^86^. A negative control was included for each culture by adding 2 mg of protein extract that was previously inactivated at 80 °C for 1 h. Tubes were incubated at 37 °C in a heating block (Thermo Fisher Scientific, Waltham, MA, USA), and the reaction was initiated by adding TMA to a final concentration of 100 mM. Aliquots of 25 µL were taken at indicated times and added to 700 µL of fresh Ellman’s reagent (0.5 mM 5,5’-dithio-*bis*-(2-nitrobenzoic acid) in 150 mM Tris pH 8.0) to stop the reaction. Color development in Ellman’s reagent was used to follow free thiols (coenzyme M) in solution^88^ and quantified at 412 nm along a coenzyme M calibration curve in a microplate reader (BioTek Epoch 2, Winooski, VT, USA) outside the anaerobic chamber. One unit was defined as the amount of enzyme that catalyzes the consumption of 1 µmol of coenzyme M per minute, and the specific activity values were calculated from 35 min reactions. All enzyme kinetics were conducted in triplicate and statistical analyses were performed with unpaired Welch *t*-tests in GraphPad Prism v10.3.1.

### Affinity purification of tagged proteins

Late-exponential phase cultures (0.5–2 L) were harvested, either aerobically or anaerobically as indicated, by centrifugation at 5,000 ×*g* for 10 mins at 4 °C (Sorvall Legend XTR, Thermo Fisher Scientific, Waltham, MA, USA) using 250 mL tight-sealed polypropylene Nalgene centrifuge bottles. The cell pellet from each 500 mL culture was lysed by resuspension in 5 mL of 50 mM Tris solution pH 8.0 containing 2 U/mL DNase I (Thermo Fisher Scientific, Waltham, MA, USA) and 1X Roche cOmplete EDTA-free Protease Inhibitor Cocktail (MilliporeSigma, Burlington, MA, USA) overnight at 4 °C. Sodium chloride was slowly added from a 5 M solution to the cell lysate with mild agitation to a final concentration of 300 mM, followed by centrifugation at 18,000 ×*g* for 60 min at 4 °C (Sorvall Legend XTR, Thermo Fisher Scientific, Waltham, MA, USA). The cleared supernatant was loaded on a 0.3–0.6 mL Strep-Tactin Superflow Plus resin (QIAGEN, Hilden, Germany) gravity-flow column equilibrated with 50 mM Tris pH 8.0, 300 mM NaCl. The column was washed seven times with 2 mL of cold 50 mM Tris pH 8.0, 300 mM NaCl, and the protein was eluted in six 0.5 mL fractions with an elution buffer comprised of 2.5 mM desthiobiotin, 50 mM Tris pH 8.0, 300 mM NaCl. The amount of protein in each fraction was monitored at 280 nm using a Nanodrop One UV-Vis Spectrophotometer (Thermo Fisher Scientific, Waltham, MA, USA) and concentrated with a 10 kDa MWCO Amicon centrifugal filter (MilliporeSigma, Burlington, MA) if required. Affinity purification and protein concentration was carried out aerobically unless otherwise noted. Protein concentration was determined using the theoretical extinction coefficient at 280 nm for tagged MA2561 (53,080 M^−1^cm^−1^) in a Nanodrop UV-Vis Spectrophotometer and by the Pierce Bradford Plus Protein Assay Reagent (Thermo Fisher Scientific, Waltham, MA, USA) against a standard curve of BSA^87^ measured in a microplate reader (BioTek Epoch 2, Winooski, VT, USA).

### Differential Scanning Fluorimetry

MA2561 was purified from cultures grown in HS media with 150 mM methanol to assess its thermal stability in the presence of different substrates via differential scanning fluorimetry. An aliquot of 0.5 mg/mL MA2561 (from strain DDN183) or 0.5 mg/mL Mcr (from strain WWM1086^89^) was incubated at room temperature for 15 min with 50 mM of the indicated substrate in a total reaction volume of 140 µL in 50 mM Tris pH 8.0, 300 mM NaCl. A fresh 1:100 dilution of SYPRO Orange Protein Gel Stain (Invitrogen, Thermo Fisher Scientific, Waltham, MA, USA) was prepared in the reaction buffer and 15.5 µL of the dilution was added to the protein-ligand binding reaction for a final SYPRO concentration of 5X. After gentle homogenization, 50 µL of the reaction mixture was dispensed in triplicate in 96-well PCR plates (Applied Biosystems, Thermo Fisher Scientific, Waltham, MA, USA) and sealed with adhesive optical film. SYPRO Orange fluorescence was measured through a temperature gradient (25–95 °C) with increments of 0.5 °C every 30 s in a CFX96 Touch Real-Time PCR Detection System (Bio-Rad, Hercules, CA) using the FRET channel. Melt peaks were plotted as the first derivative of the fluorescence emission as a function of temperature (dF/dT), and the protein melting temperature (T_m_)—defined as the midpoint temperature in the transition to maximum fluorescence and interpreted as the temperature at which 50% of the protein unfolds—was calculated from a quadratic curve fit to the closest 5 datapoints in the derivative peak. Statistical difference among the T_m_ obtained from triplicate melt curves in the presence of different ligands was evaluated in GraphPad Prism v10.3.1 using unpaired *t*-tests with Welch’s correction.

### SDS-PAGE and Western Blot

Protein concentrates, fractions, or cell lysates were mixed with Laemmli sample buffer^90^ under reducing conditions (2.5% β-mercaptoethanol) and boiled for 5 mins at 100 °C, unless otherwise indicated. Samples were loaded on 4–20% gradient or 12% precast Tris-Glycine denaturing SDS-PAGE gels (Mini-PROTEAN TGX, Bio-Rad, Hercules, CA, USA) along with the Bio-Rad Precision Plus Prestained Protein Standards, or the Invitrogen PeppermintStick Phosphoprotein Molecular Weight Standards (Thermo Fisher Scientific, Waltham, MA, USA), and run at 80–120V under aerobic conditions unless otherwise noted. SDS-PAGE gels were stained with Coomassie GelCode Blue Stain Reagent or with Pro-Q Diamond Phosphoprotein Gel Stain (Thermo Fisher Scientific, Waltham, MA, USA) according to the manufacturer’s instructions. Mass Spectrometry sample prep and Protein ID were performed by Applied Biomics, Inc (Hayward, CA, https://www.appliedbiomics.com/). Samples that were visualized by immunostaining were transferred from the gel to a PVDF membrane using the Trans-Blot Turbo Transfer System and Midi PVDF Transfer Pack (Bio-Rad, Hercules, CA) as per the manufacturer’s instructions. The PVDF membranes were rinsed with TBS-T (20 mM Tris–HCl pH 7.6, 150 mM NaCl, 0.1% Tween-20), blocked for 1 h with 5% nonfat milk dissolved in TBS-T, and incubated at room temperature for 60 mins with mouse monoclonal anti-FLAG M2 HRP-conjugated antibody (Sigma-Aldrich, St Louis, MO) diluted 1:100,000 in TBS-T. After 6 final washes of 5 min with TBS-T, the signal was developed by incubating the membranes for 5 min in the dark with the Immobilon Western Chemiluminescent HRP Substrate (MilliporeSigma, Burlington, MA). Gels and blots were visualized on a ChemiDoc MP Imaging System (Bio-Rad, Hercules, CA).

### Quantification of free thiols

Exposed sulfhydryl groups (or free thiols) were quantified by Ellman’s reagent and used to estimate the number of reduced cysteines in MA2561. Purified MA2561 from cultures grown in HS media with 150 mM methanol was dispensed in 96-well microplates at a concentration range of 0.78–2 mg/mL in 50 µL 50 mM Tris pH 8.0, 300 mM NaCl. A calibration curve with cysteine (0–250 µM; bioWORLD, Dublin, OH, USA) dissolved in the reaction buffer, along with 50 µL of 2 mg/mL BSA (Pierce Albumin Standard, Thermo Fisher Scientific, Waltham, MA, USA) was included in the microplate. All reactions for proteins at each concentration and for cysteine standards were conducted in triplicate. Ellman’s reagent^88^ was prepared by making a 0.5 mM 5,5’-dithio-*bis*-(2-nitrobenzoic acid) solution in 150 mM Tris pH 8.0. A total of 200 µL of fresh Ellman’s solution was added to each well of the microtiter plate and incubated for 10 min at room temperature. Absorbance was measured at 412 nm in a microplate reader (BioTek Epoch 2, Winooski, VT, USA) and the number of free thiols per protein was calculated relative to the cysteine standard curve. This assay was performed aerobically or anaerobically (inside an anaerobic chamber with a 4:96 H_2_:N_2_ atmosphere) as indicated, however, all absorbance readings were carried out outside the anaerobic chamber, as the color of 2-nitro-5-thiobenzoate in the reaction mixture appeared stable in the presence of oxygen.

### Electrophoretic Mobility Shift Assay (EMSA)

A 5’ Cy5-labeled primer (Integrated DNA Technologies, Coralville, IA, USA) targeting a region upstream of the *mtmCB1* promoter was used to generate fluorescently labeled oligonucleotide probes of different lengths covering the promoter and 5’-UTR sequence. Probes were purified from agarose gels (Zymoclean Gel DNA Recovery kit, Zymo Research, Irvine, CA, USA). For the EMSA, 5 nM DNA was incubated with MA2561 (0.04–1.3 mg/mL) for 30 min in the dark at room temperature in 30 µL of binding reaction containing 50 mM Tris pH 8.0, 300 mM NaCl, 0.1% v/v Nonidet P-40, and 10% v/v glycerol. Binding controls were incubated in the presence of 1.3 mg/mL MA2561 with the addition of 2.5 µM cold probe, or with 2.5% β-mercaptoethanol (β-ME). Additional controls using 1.3 mg/mL BSA, or 1.3 mg/mL of other purified proteins from *M. acetivorans* (Mcr^89^, MA2555, and MA4396) were included. The entire reaction volume was loaded on a fresh native 4% polyacrylamide (60:1 acrylamide:bisacrylamide) gel prepared with 0.5X TBE buffer (45 mM Tris-borate pH 8.3, 1 mM EDTA) and 0.1% Nonidet P-40 cast in 19.5×16 cm glass plates with 1 mm thick spacers. Gels were run for 5 h at 40 V at room temperature in the dark with 0.5X TBE running buffer in a vertical GibcoBRL V16 tank under aerobic conditions, unless otherwise noted. For gel shifts in the presence of methanogenesis substrates, 10 mM TMA, DMA, MMA, or methanol were added to the binding reaction, gel, and running buffer. Gels were visualized using an Amersham Typhoon Trio+ imager with a 633 nm laser and a 670 nm emission filter (GE Healthcare, Chicago, IL, USA), unless otherwise indicated. Band intensity was quantified from the raw images using ImageJ2 v2.14.0/1.54f, and the levels of the images in the figures were uniformly adjusted to ease visualization.

### Phylogenetic analyses

The 16S rRNA gene phylogeny of representative methanogens within the class *Methanosarcinia* were used to evaluate the phyletic distribution of AmzR. The sequences of 16S rRNA genes were obtained from the NCBI nucleotide collection or the IMG database, aligned with MAFFT v7.309^91^, and maximum likelihood trees were built with PhyML v3.2^92^ using 1,000 bootstraps in Geneious v9.1.8. Three *Methanocella* spp. (*M. paludicola* SANAE, *M. arvoryzae* MRE50, and *M. conradii* HZ254) were included in the analysis as closely related outgroups of *Methanosarcinia*. For the AmzR phylogenetic tree, sequences were retrieved with BLASTp v2.15.0^93^ using MA2561 as query. Only the closest MA2561 protein homolog with the lowest E-value in each genome was used for building the maximum likelihood phylogenetic tree as described above. Sequence logo of AmzR was obtained from a MAFFT v7.309 alignment in Geneious v9.1.8, and tanglegrams comparing the 16S rRNA gene and AmzR trees were built in Dendroscope v3.8.10. The presence of AmzR and methylamine methyltransferases coded in each genome was determined with BLASTp v2.15.0^93^ using a E-value cutoff of 10^-50^ and 40% sequence identity in the NCBI nr database against the *M. acetivorans* MA2561, MtmC1, MtbC1, and MttC1 protein sequences (NCBI Reference Sequence WP_011022527.1, WP_011020203.1, WP_011020576.1, and WP_011020578.1, respectively). The environmental distribution of each organism was determined by searching their sample abundances of each organism in MicrobeAtlas v1.0^94^ or based on the source noted for representative isolates in NCBI. Correlation between the presence of *amzR* and *mtmC1*, *mtbC1*, or *mttC1* was determined by a Fisher’s exact test of data from genomes with annotated proteins in the class *Methanosarcinia* (>90% completeness by CheckM) that are listed as representative species in the Genome Taxonomy Database (GTDB) release 220^95^. Gene neighborhood comparison was visualized using clinker & clustermap^96^.

### Protein structural predictions

AlphaFold 3^54^ was used to obtain structural models of dimeric *M. acetivorans* C2A MA2561, *M. mazei* Gö1 MM_3108, *M. barkeri* MS MSBRM_1623 and MSBRM_1624 (NCBI Reference Sequence WP_011022527.1, WP_048037404.1, WP_048117785.1, and WP_048117783.1, respectively), and dimeric *Halobacterium salinarum* GvpE (NCBI Reference Sequence WP_010904106.1) complexed with DNA containing the GvpE-responsive element sequence^97,98^. Structural models were visualized using ChimeraX v1.8^99^. Protein domains were identified using InterPro v102.0^100^.

## Acknowledgements

We thank all members of the Nayak lab for their valuable feedback and support. We would like to specifically acknowledge Annelise Goldman for helpful discussions and technical support. We would also like to acknowledge Donald Rio for technical advice with gel shift analyses. FMF is supported by the Simons Postdoctoral Fellowship in Marine Microbial Ecology. DDN acknowledges funding from the Searle Scholars Program sponsored by the Kinship Foundation, the Rose Hills Innovator Grant, the Beckman Young Investigator Award sponsored by the Arnold and Mabel Beckman Foundation, the Alfred P. Sloan Research Fellowship sponsored by the Sloan Foundation, the Simons Foundation Early Career Investigator in Marine Microbial Ecology and Evolution Award, and the Packard Fellowship in Science and Engineering sponsored by the David and Lucille Packard Foundation. DDN is a Chan-Zuckerberg Biohub – San Francisco Investigator. The funders had no role in the conceptualization and writing of this manuscript or the decision to submit the work for publication.

## Author contributions

FMF participated in conceptualization, data curation, formal analysis, investigation, methodology, supervision, validation, visualization, writing – original draft preparation, and writing – review and editing. DDN was involved in conceptualization, funding acquisition, investigation, methodology, project administration, resources, supervision, validation, writing – original draft preparation, and writing – review and editing.

## Resource availability

Sequencing data have been deposited in the Sequencing Reads Archive and the Bioproject number will be made available upon publication. All other data generated in this study are provided in the manuscript.

## Supplemental information

Document S1. Figures S1–S27

Document S2. Excel file containing Tables S1–S11.

