## Supplement for "An Archaeal Transcription Factor Bridges Prokaryotic and Eukaryotic Regulatory Paradigms"

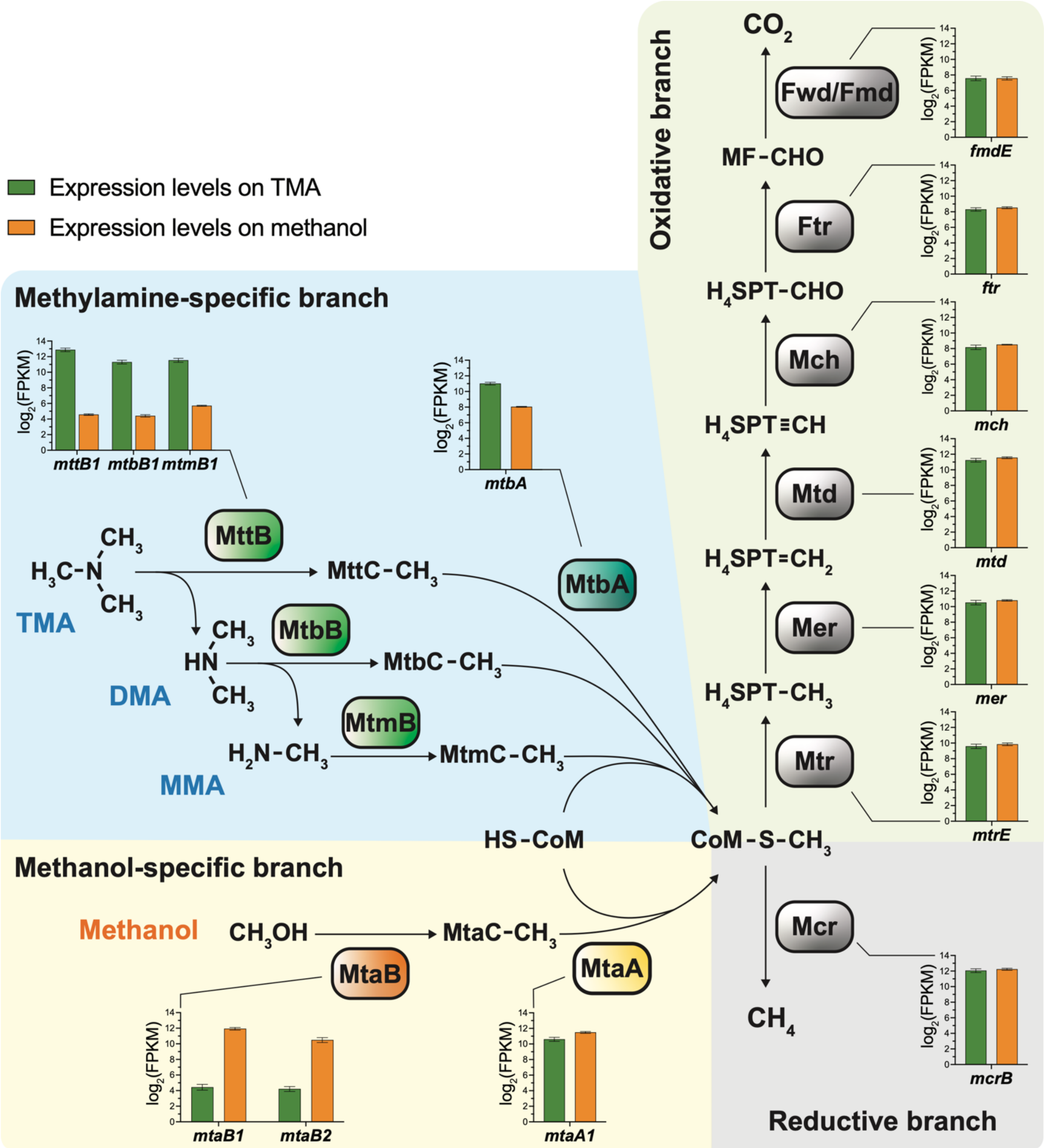


**Figure S1.** Methylotrophic methanogenesis in *Methanosarcina acetivorans* (related to Figure 1A). During growth on trimethylamine (TMA), methylamine-specific methyltransferases MttB, MtbB, and MtmB catalyze the transfer of one methyl group from TMA, dimethylamine (DMA), and monomethylamine (MMA) to the corrinoid-binding protein MttC, MtbC, and MtmC, respectively. The complex between the methyltransferase and the corresponding corrinoid protein is referred as the Methyltransferase 1 (MT1). The methyl group from the corrinoid protein is then transferred to coenzyme M (HS-CoM) by MtbA, forming methyl coenzyme M ($CH_3$-CoM). Analogous MT1 and MT2 reactions are utilized during growth on methanol via the methanol-specific MtaB, MtaC, and MtaA proteins respectively. The methyl group in $CH_3$-CoM is either reduced to methane or oxidized to carbon dioxide in a 3:1 ratio. While the enzymes in the reductive and oxidative branches (gray) remain constitutively expressed, the methylamine-specific methyltransferases (green), particularly the MT1s, are significantly upregulated in media containing TMA. Similarly, methanol-specific MT1s (orange) are upregulated only in media with methanol. Graphs indicate expression values of genes encoding each enzyme of the pathway as FPKM (fragments per kilobase of transcript per million mapped reads). Only the expression level of one representative subunit is represented for multi-subunit enzymes. Expression values are only shown for isoforms of the MT1 and MT2 that have been experimentally validated.

Mcr, methyl-coenzyme M reductase; Mtr, methyl-tetrahydrosarcinopterin:coenzyme M methyltransferase; Mer, methylene-tetrahydrosarcinopterin reductase; Mtd, methylene-tetrahydrosarcinopterin dehydrogenase; Mch, methenyl-tetrahydrosarcinopterin cyclohydrolase; Ftr, formylmethanofuran:tetrahydrosarcinopterin formyl-transferase; Fwd/Fmd, formylmethanofuran dehydrogenase.

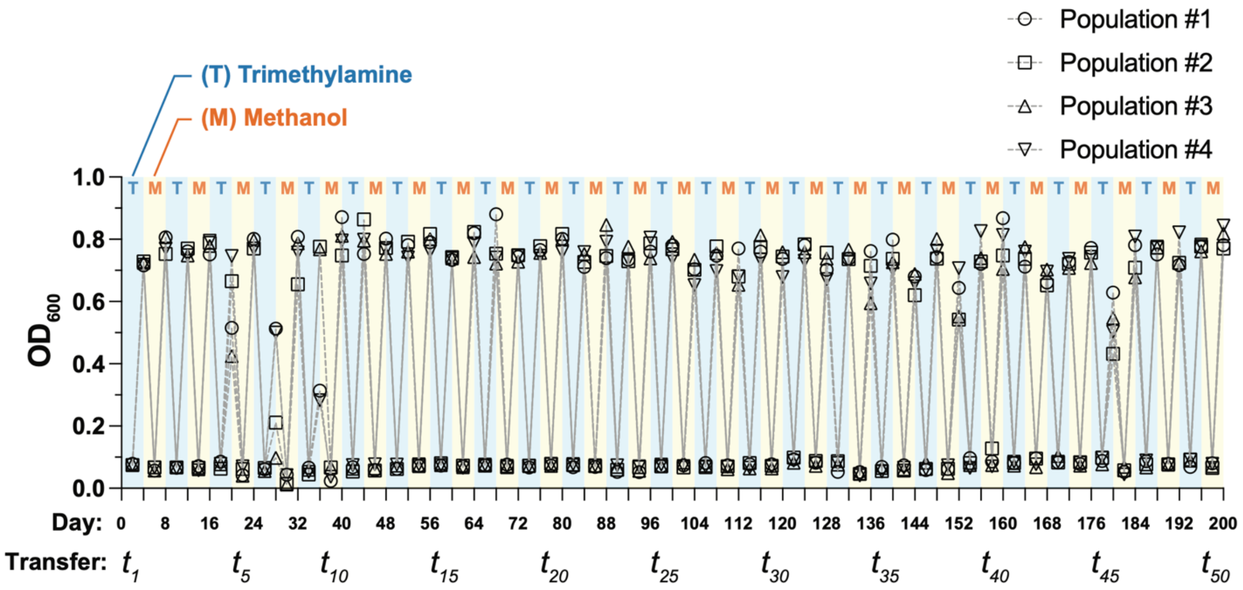


**Figure S2.** Initial and final $OD_{600}$ of four replicate populations of *M. acetivorans* WWM60 for the duration of the evolution experiment (shown in Figure 1C). Cells were alternately grown in minimal media containing either 25 mM trimethylamine (T) or 75 mM methanol (M) as the sole methanogenic substrate. The alternating substrate regime was repeated for a total of fifty transfers ($t_1$–$t_{50}$) every 4 days using a 1:20 dilution of the previous passage. The evolution experiment lasted 200 days or *ca.* 200 generations.

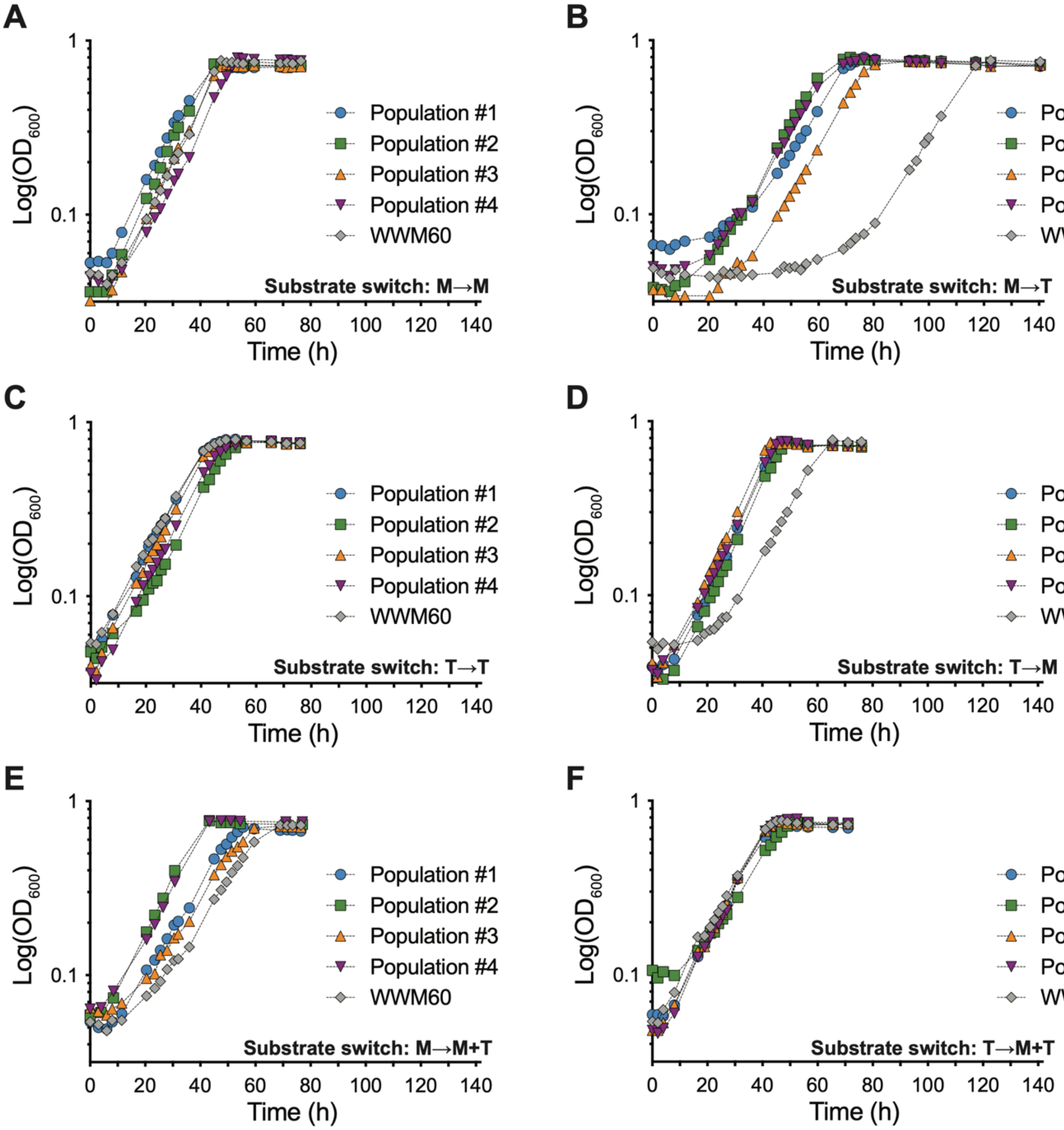


**Figure S3.** Phenotypic evaluation of evolved *M. acetivorans* populations. Growth curves of the parent strain (WWM60) and the evolved Populations #1–4 ($t_{50}$; see Figure 1C and Figure S2) after methanol-grown cells are transferred to media with (**A**) methanol (M→M) or **(B)** TMA (M→T), TMA-grown cells are transferred to media with (**C**) TMA (T→T) or (**D**) methanol (T→M), (**E**) methanol-grown cells are transferred to media with methanol and TMA (M→M+T), and (**F**) TMA-grown cells are transferred to media with methanol and TMA (T→M+T).

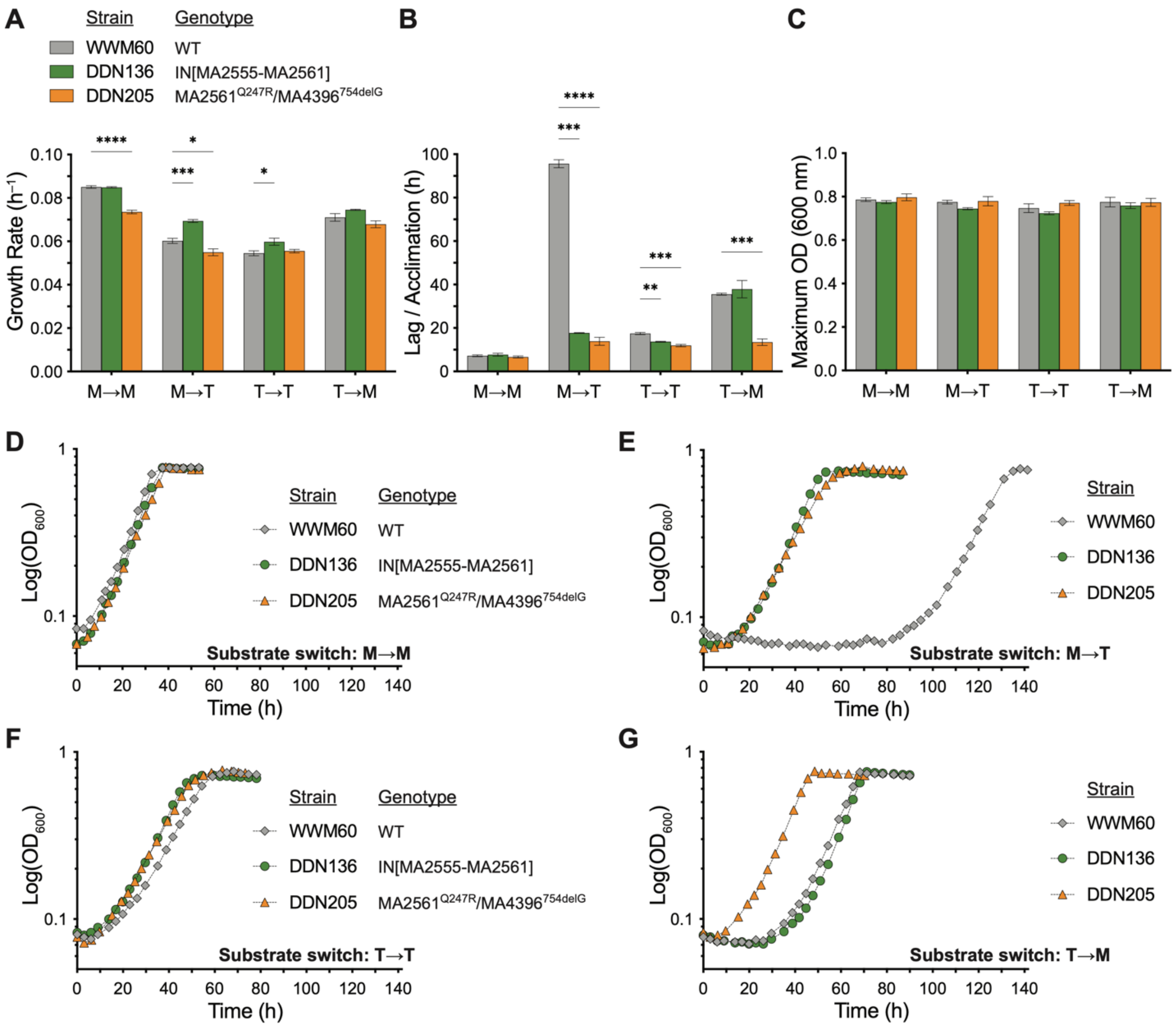


**Figure S4.** Growth parameters of isolates from the evolved populations (see Figure 1D). (**A**) Mean growth rate, (**B**) lag/acclimation time, and (**C**) growth yield of the parent strain (WWM60; gray), an isolate from Population #1 (DDN136, containing the genomic inversion IN[MA2555-MA2561]; green) and an isolate from Population #4 (DDN205, containing the mutations MA2561$^{Q247R}$ and MA4396$^{754delG}$; orange). All growth experiments were conducted in triplicate. Error bars represent one standard deviation (SD) and growth parameters of isolates were compared to WWM60 using unpaired *t*-tests with Welch correction (*P<0.05, **P<0.01, ***P<0.001, ****P<0.0001). (**D**) Semilogarithmic plot of representative growth curves of WWM60 (WT; gray), DDN136 (green), DDN205 (orange) when methanol-grown cells are transferred to methanol-containing media (M→M), (**E**) methanol-grown cells are transferred to TMA-containing media (M→T; plot shown in Figure 1D but repeated here for comparison with other growth conditions), (**F**) TMA-grown cells are transferred to TMA-containing media (T→T), or (G) TMA-grown cells are transferred to methanol-containing media (T→M).

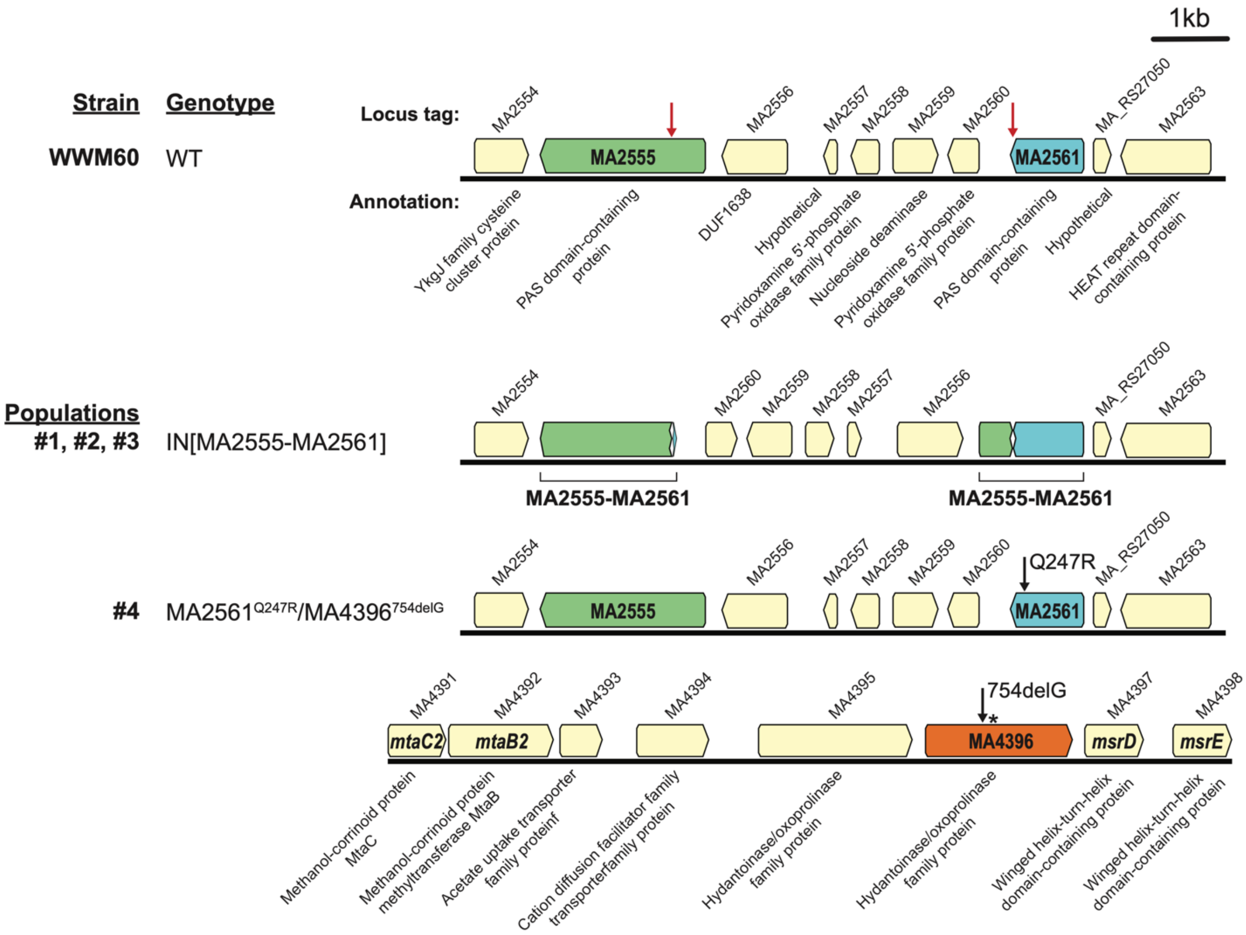


**Figure S5.** Mutations fixed in four evolved populations ($t_{50}$; related to Figures 1C and 1D). Populations #1, #2, and #3 have the same genomic inversion between two insertion sites located within the coding sequence of MA2555 and MA2561 (red arrows in WWM60). The genomic inversion, IN[MA2555-MA2561], results in premature truncation of MA2555 and an extension of MA2561 after R314 by 106 amino acids. Population #4 has a point mutation in MA2561 (Q247R) and a 1-bp frameshift deletion (754delG) in the MA4396 locus, which are indicated by black arrows. DDN136 is an isolate from population #1 with the IN[MA2555-MA2561] mutation, and DDN205 is an isolate from population #4 with MA2561$^{Q247R}$/MA4396$^{754delG}$ mutations. These isolates were used for in-depth phenotypic characterization of the genotypes as shown in Figure 1D and S4.

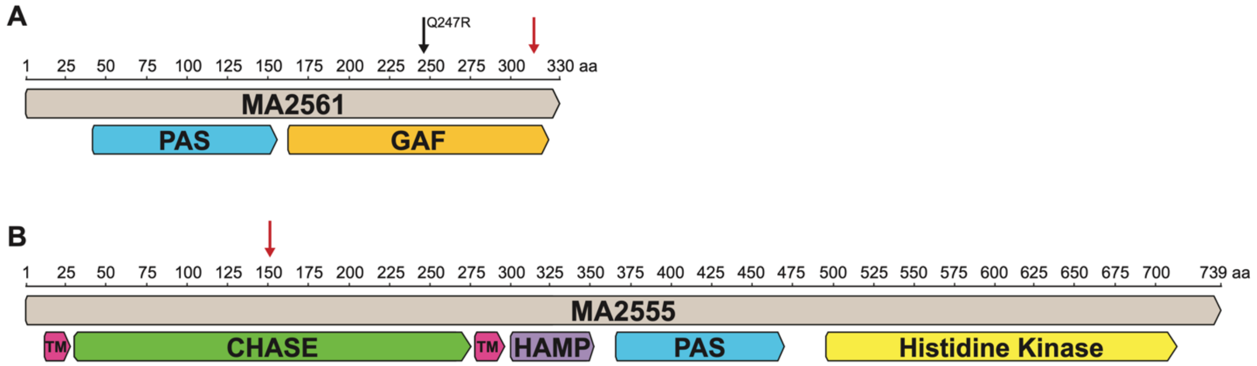


**Figure S6.** MA2561 and MA2555 have domains commonly found in regulatory proteins. (**A**) Sequence representation of MA2561, a predicted cytosolic protein of 330 amino acids containing one PAS (blue) and one GAF (orange) domain. Black and red arrows show the position of the point mutation and the inversion site, respectively, after the experimental evolution (see Figure S5). (**B**) MA2555 is predicted to be an integral membrane protein with two putative transmembrane domains (TM) (pink), an extracellular CHASE domain (green), and intracellular HAMP (purple), PAS (blue), and Histidine Kinase (yellow) domains. Red arrow shows the insertion site for the genomic inversion after the experimental evolution (see Figure S5). All protein domains were identified using InterPro 102.0[1].

PAS: Per-Arnt-Sim domain

GAF: cGMP-specific phosphodiesterases, Adenylyl cyclases and FhlA domain

CHASE: Cyclases/Histidine kinases Associated Sensory Extracellular domain

GAF: cGMP-specific phosphodiesterases, Adenylyl cyclases and FhlA domain

HAMP: Histidine kinases, Adenylate cyclases, Methyl accepting proteins and Phosphatases domain

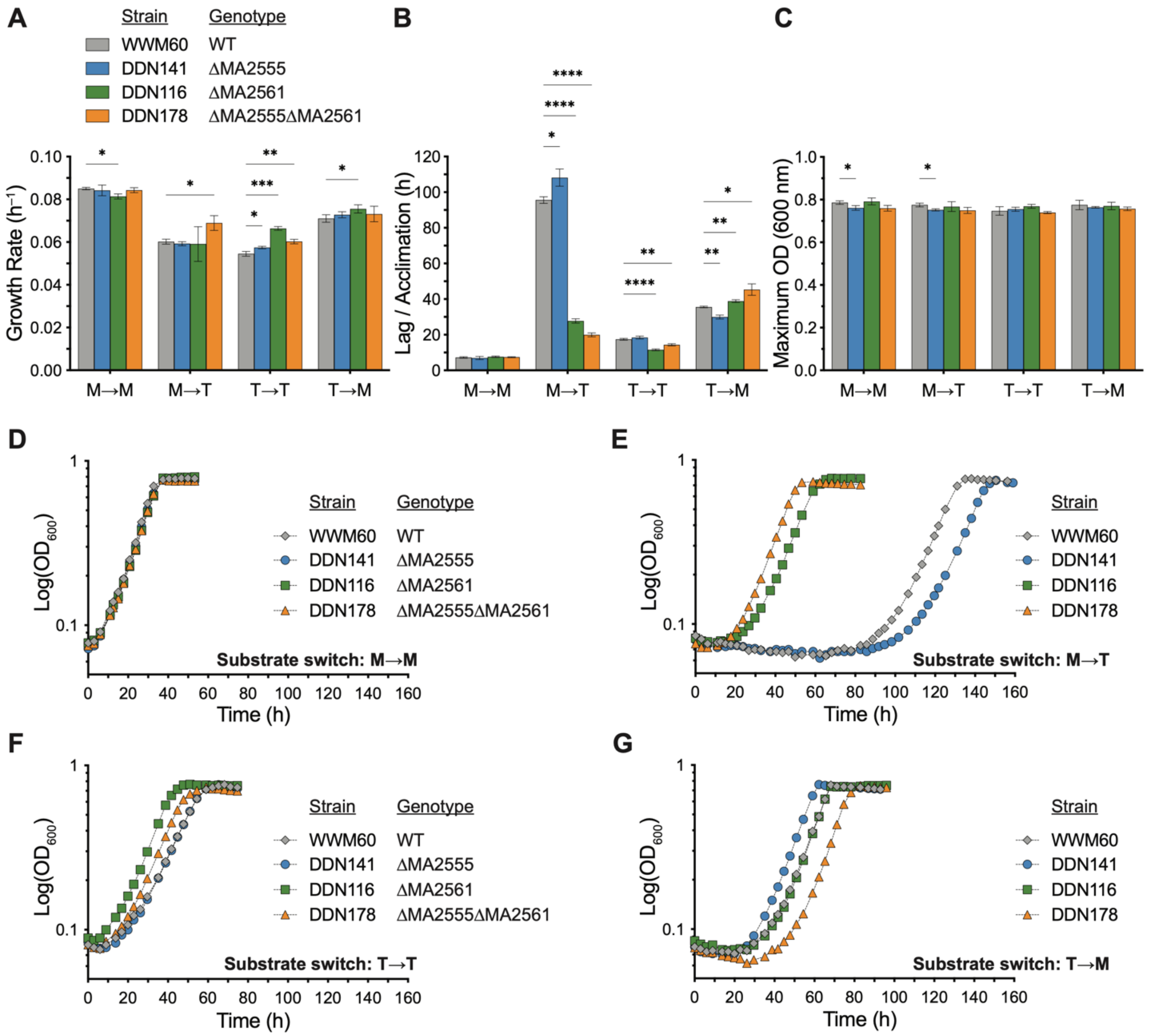


**Figure S7.** Phenotypic characterization of ΔMA2555, ΔMA2561 and ΔMA2555ΔMA2561 mutants on a single substrate (related to Figure 1E). (**A**) Growth rate, (**B**) lag/acclimation time, and (**C**) growth yield of the parent strain (WWM60; gray), the ΔMA2555 mutant (DDN141; blue), the ΔMA2561 mutant (DDN116; green) and the ΔMA2555ΔMA2561 double-knockout mutant (DDN178; orange). All growth experiments were conducted in triplicate. Error bars represent one standard deviation (SD) and growth parameters of isolates were compared to WWM60 using unpaired *t*-tests with Welch correction (*P<0.05, **P<0.01, ***P<0.001, ****P<0.0001). (**D**) Representative growth curves of WWM60 (WT; gray), ΔMA2555 (DDN141; blue), ΔMA2561 (DDN116; green) and ΔMA2555ΔMA2561 (DDN178; orange) when methanol-grown cells are transferred to methanol-containing media (M→M), (**E**) methanol-grown cells are transferred to TMA-containing media (M→T; data shown in Figure 1E and repeated here for comparison), (**F**) TMA-grown cells are transferred to TMA-containing media (T→T), or (**G**) TMA-grown cells are transferred to methanol-containing media (T→M). Only strains with deletions in MA2561 show faster M→T acclimation times.

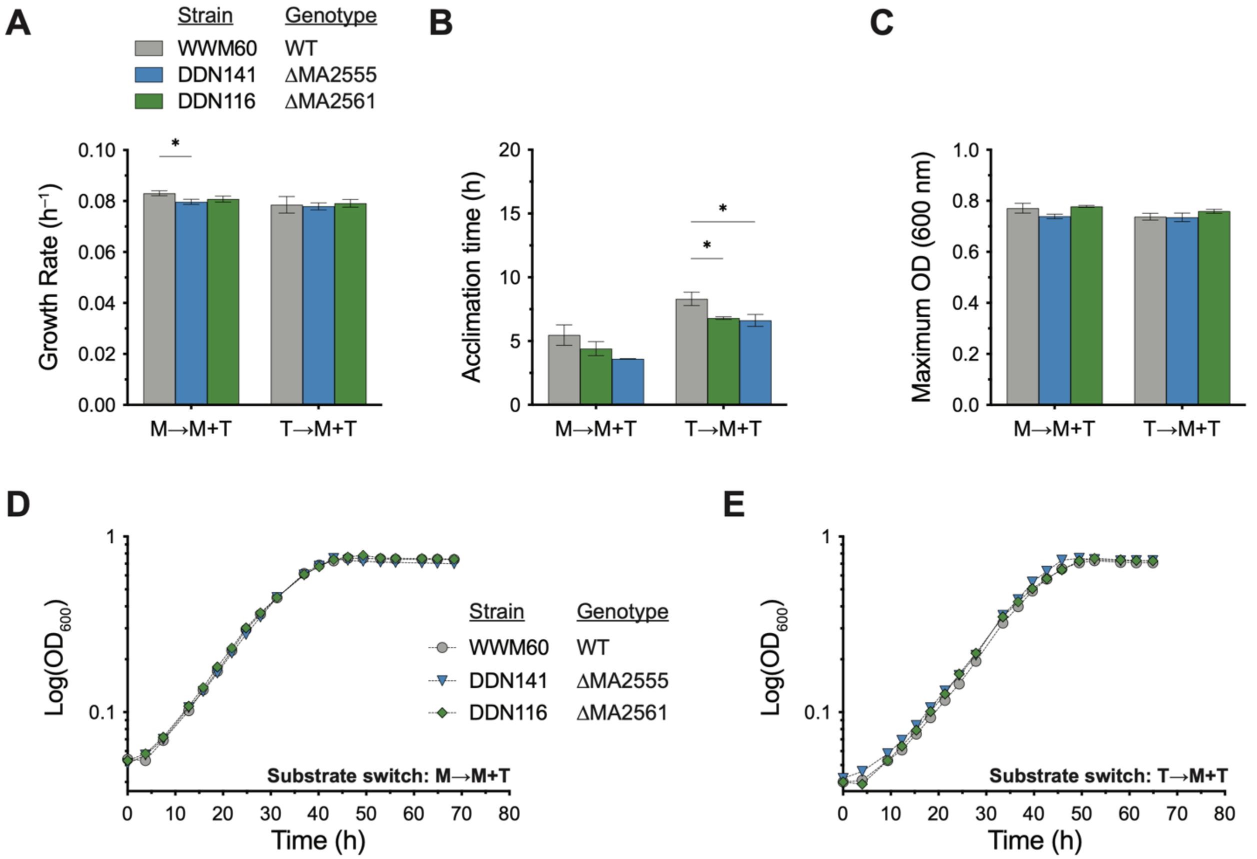


**Figure S8.** Phenotypic characterization of ΔMA2555, ΔMA2561 and ΔMA2555ΔMA2561 mutants in dual-substrate media (related to Figure 1E). (**A**) Growth rate, (**B**) acclimation time, and (**C**) growth yield of the parent strain (WWM60; gray), the ΔMA2555 mutant (DDN141; blue) and the ΔMA2561 mutant (DDN116; green) when cells growth on either methanol (M→M+T) or TMA (T→M+T) were transferred to medium containing both methanol and TMA. All growth experiments were conducted in triplicate. Error bars represent one standard deviation (SD) and growth parameters of isolates were compared to WWM60 using unpaired *t*-tests with Welch correction (*P<0.05, **P<0.01, ***P<0.001, ****P<0.0001). Representative growth curves in media containing both methanol and TMA (M+T) after transfer from either a (**D**) methanol-grown inoculum, or a (**E**) TMA-grown inoculum are shown.

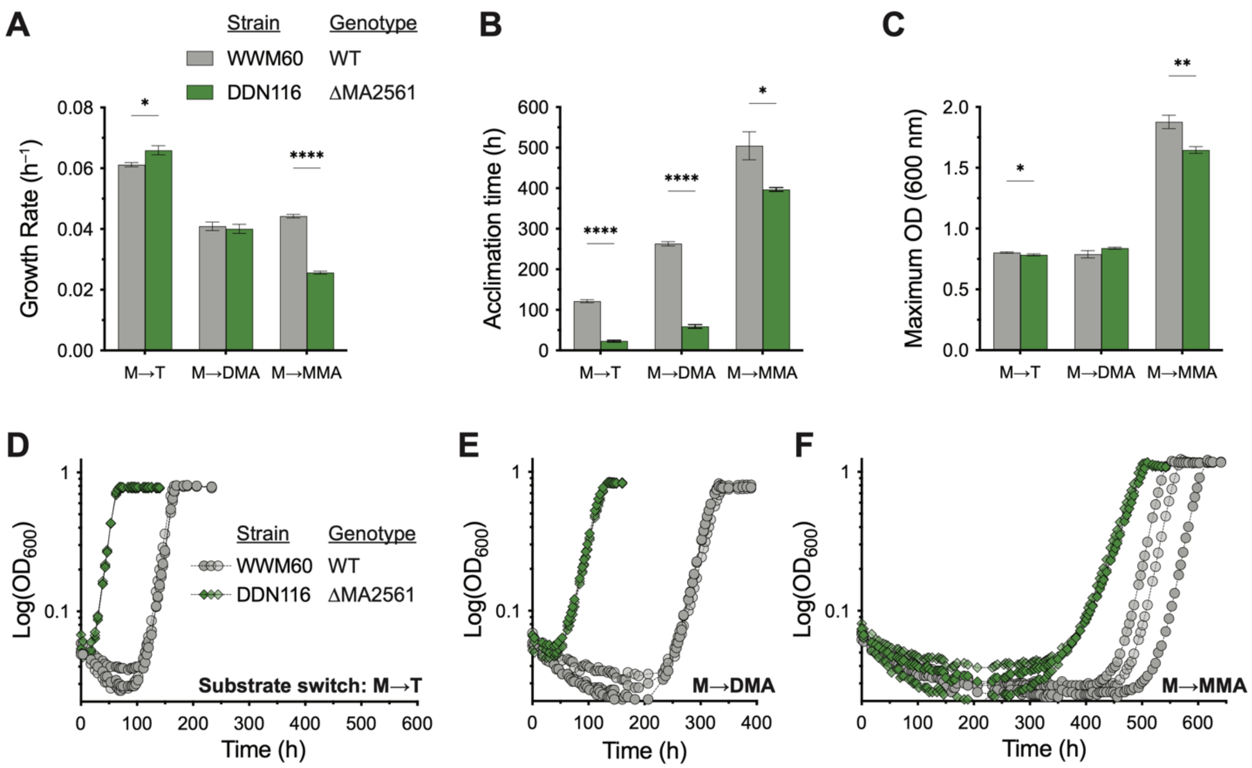


**Figure S9.** Phenotypic characterization of ΔMA2561 mutant on methylamines (related to Figure 1F and 1G). (**A**) Growth rate, (**B**) acclimation time, and (**C**) growth yield of the parent strain (WWM60; gray) and the ΔMA2561 mutant (DDN116; green) on trimethylamine (T), dimethylamine (DMA) and methylamine (MMA). All growth experiments were conducted in triplicate. Error bars represent one standard deviation (SD) and growth parameters of isolates were compared to WWM60 using unpaired *t*-tests with Welch correction (*P<0.05, **P<0.01, ***P<0.001, ****P<0.0001). Triplicate growth curves of the parent strain (WWM60; gray) and the ΔMA2561 mutant (DDN116; green) after a (**D**) methanol to TMA (M→T), (**E**) methanol to DMA (M→DMA), or (**F**) methanol to MMA (M→MMA) switch are shown. Data from panels E and F are shown in Figure 1F and 1G, respectively, but repeated here for comparison.

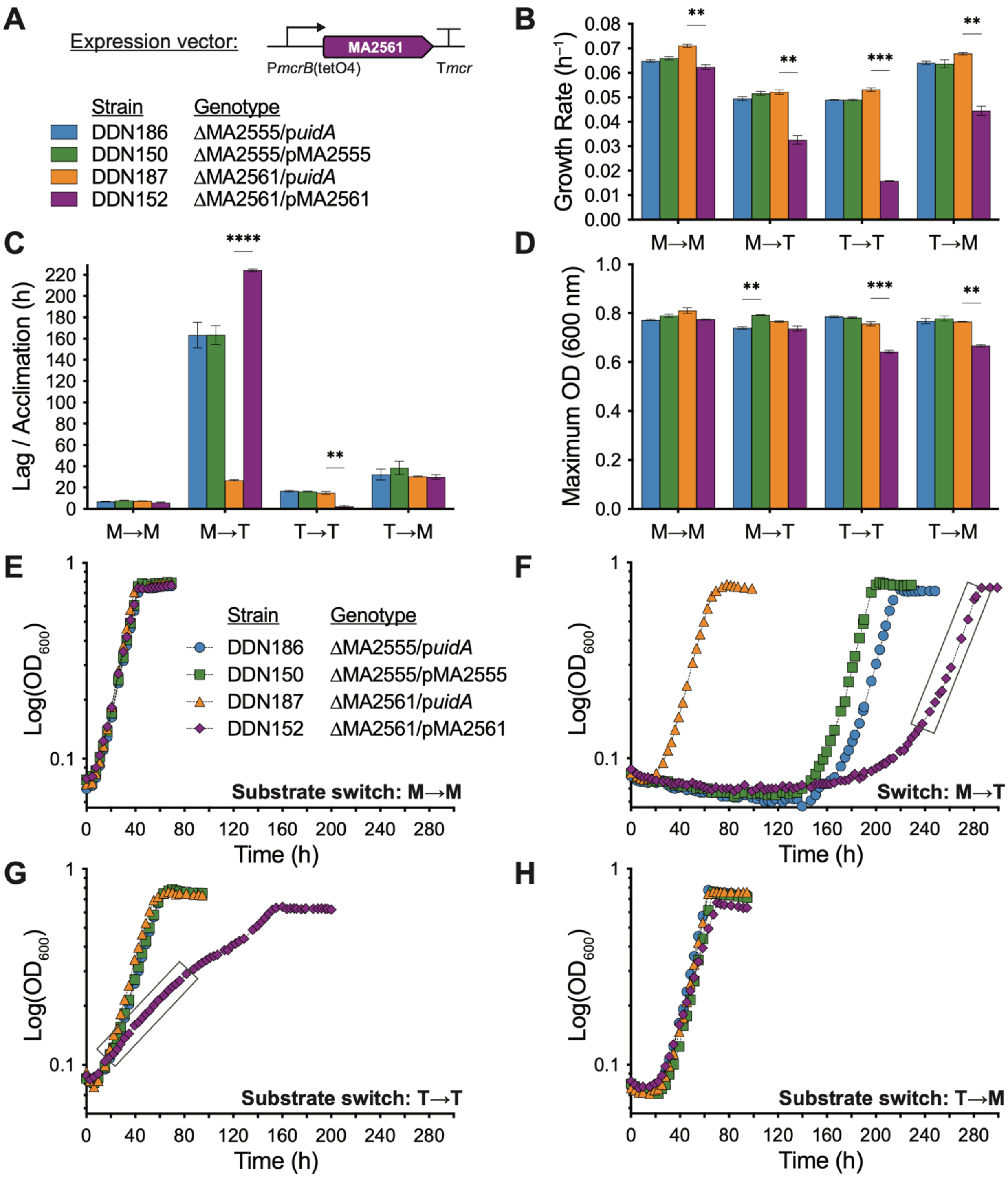


**Figure S10.** Phenotypic analyses of MA2561 complementation strains. (**A**) Schematic of the expression vector used for complementation of MA2555 or MA2561 *in trans* under the control of the tetracycline-inducible promoter P*mcrB*(tetO4). ΔMA2555 and ΔMA2561 strains were complemented with a plasmid containing the deleted gene ('pMA2555' or 'pMA2561' in DDN150 or DDN152, respectively), or a negative control plasmid containing the *E. coli* β-glucuronidase gene ('p*uidA*', DDN186 and DDN187 respectively) downstream of P*mcrB*(tetO4). (**B**) Growth rate, (**C**) lag/acclimation time, and (**D**) growth yield of DDN186 (ΔMA2555/p*uidA*; blue),

DDN150 (ΔMA2555/pMA2555; green), DDN187 (ΔMA2561/ p*uidA*; orange) and DDN152 (ΔMA2561/p*uidA*; purple). Expression from P*mcrB*(tetO4) was fully induced by the addition of 100 µg/mL tetracycline to the growth medium. All growth experiments were conducted in triplicate. Error bars represent one standard deviation (SD) and growth parameters of isolates were compared using unpaired *t*-tests with Welch correction (*P<0.05, **P<0.01, ***P<0.001, ****P<0.0001). Representative growth curves of strains overexpressing MA2555, MA2561, or UidA when (**E**) methanol-grown cells are transferred to methanol-containing media (M→M), (**F**) methanol-grown cells are transferred to TMA-containing media (M→T), (**G**) TMA-grown cells are transferred to TMA-containing media (T→T), or (**H**) TMA-grown cells are transferred to methanol-containing media (T→M). As overexpression of MA2561 led to non-exponential growth on TMA, data points that were a best fit for exponential growth in (**F**) and (**G**) are indicated by a rectangle and were used to estimate growth rates.

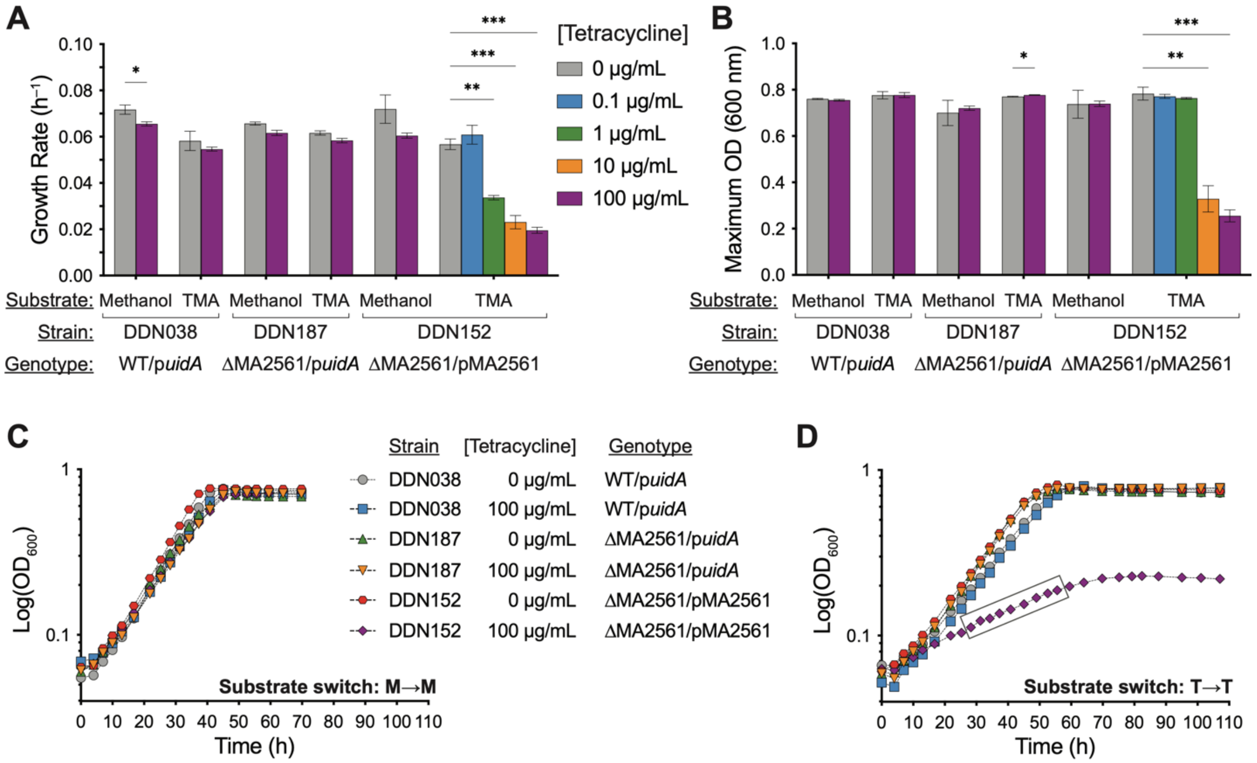


**Figure S11.** Dose-dependent inhibition of TMA growth by MA2561 (related to Figure 2A). (**A**) Growth rate and (**B**) yield on methanol or TMA of strains with a plasmid expressing either MA2561 ('pMA2561') or a control gene ('p*uidA*') under the tetracycline-inducible P*mcrB*(tetO4). Different concentrations of tetracycline were added to the culture media to tune the expression of MA2561 or *uidA*. All growth experiments were conducted in triplicate. Error bars represent one standard deviation (SD), and comparisons between cultures with (purple) and without (gray) tetracycline were conducted using unpaired *t*-tests with Welch correction (*P<0.05, **P<0.01, ***P<0.001, ****P<0.0001). (**C**) Representative growth curves on methanol or (**D**) TMA of MA2561- or *uidA*-complemented strains in the absence or presence of 100 µg/mL tetracycline. Additional growth curves from cultures with intermediate tetracycline concentrations are shown in Figure 2A. Data points used to calculate the growth rate in cells overexpressing MA2561 on TMA are enclosed in a rectangle.

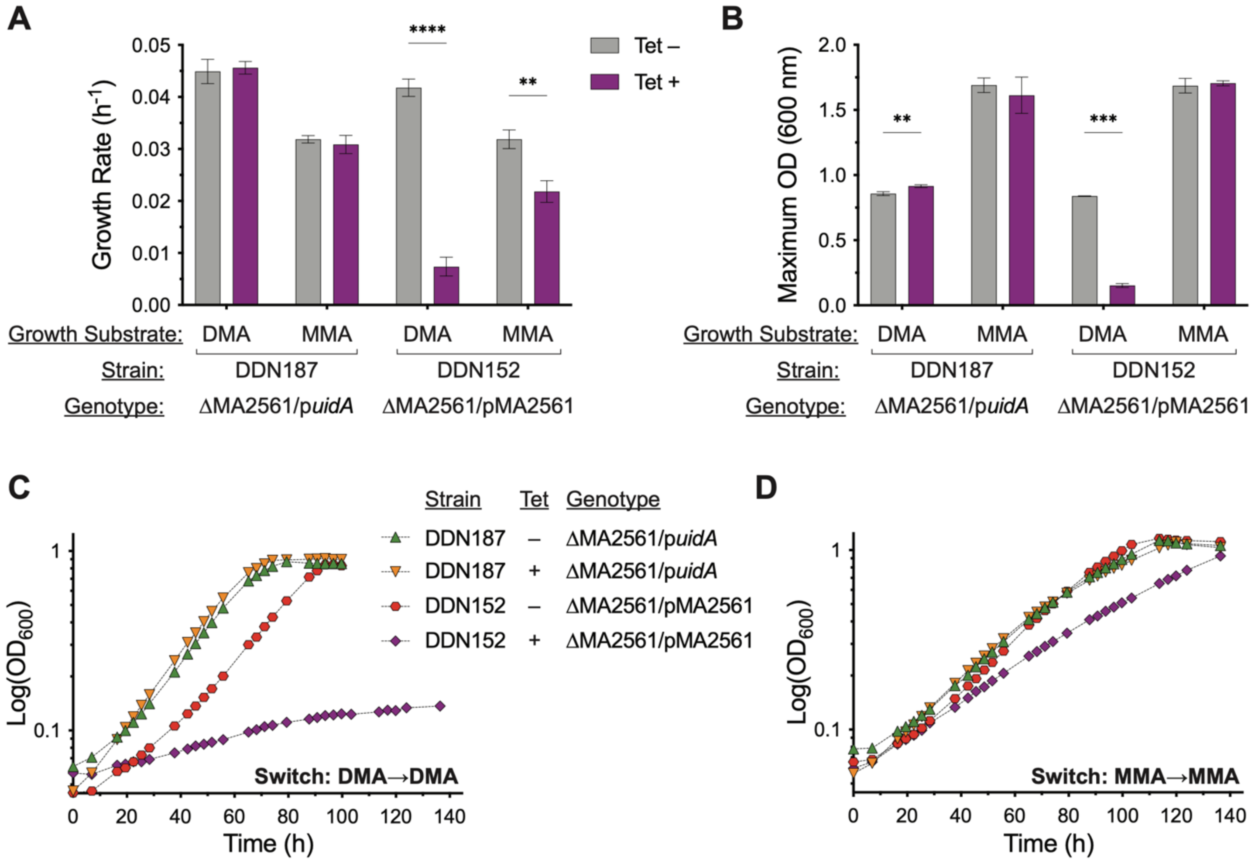


**Figure S12.** MA2561 overexpression impairs growth on methylamines (related to Figures 2B and 2C). (**A**) Growth rate and (**B**) yield of ΔMA2561 complemented with plasmids containing either *uidA* ('p*uidA*', negative control) or MA2561 ('pMA2561') under the control of the tetracycline-inducible promoter P*mcrB*(tetO4) on dimethylamine (DMA) and monomethylamine (MMA). All growth experiments were conducted in triplicate. Error bars show one standard deviation (SD), and significant differences between cultures with (purple) and without (gray) 100 µg/mL tetracycline are shown (unpaired *t*-tests with Welch correction, *P<0.05, **P<0.01, ***P<0.001, ****P<0.0001). Representative growth curves on (**C**) DMA or (**D**) MMA of ΔMA2561 containing an expression vector for either *uidA* (DDN187) or MA2561 (DDN152) in media without (–) or with (+) 100 µg/mL tetracycline (Tet). Triplicate growth curves of DDN152 are shown in Figure 2B, C.

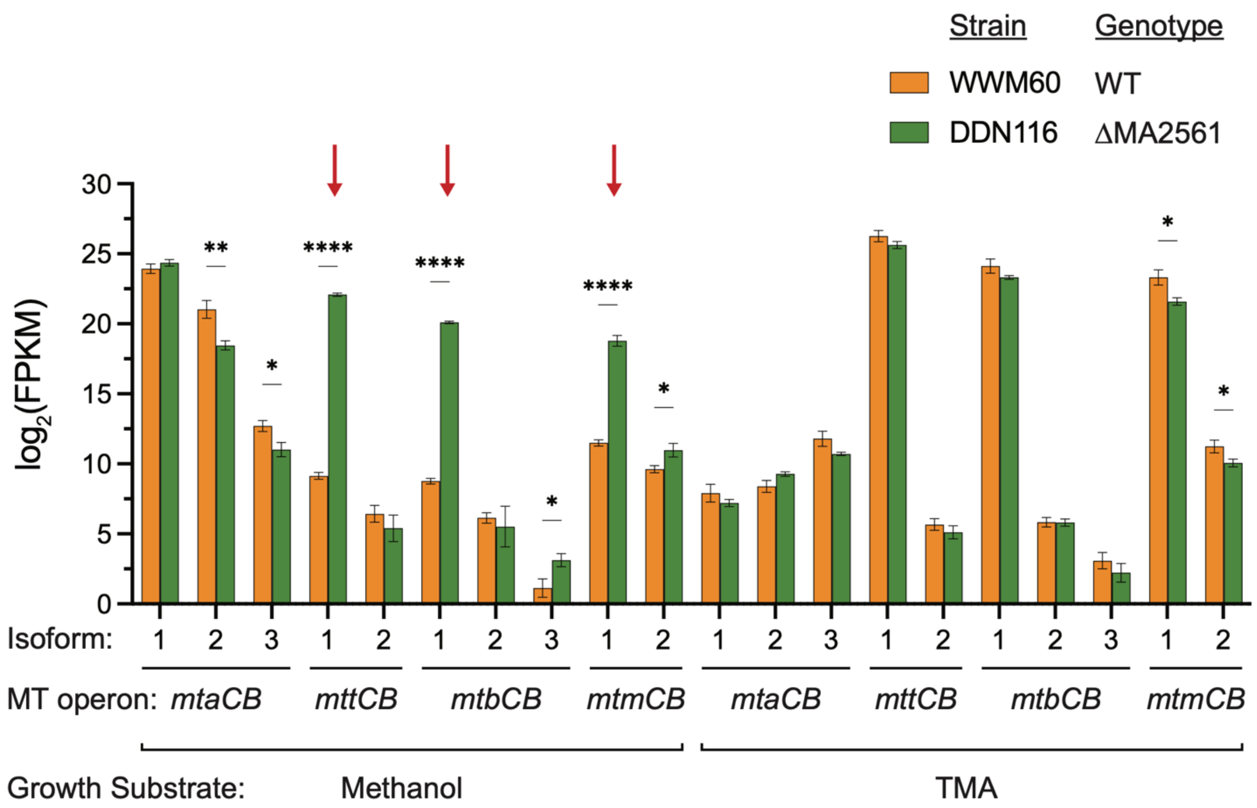


**Figure S13.** Methylamine-specific MT1s are highly expressed on methanol in the ΔMA2561 mutant (related to Figure 2D). FPKM (fragments per kilobase of transcript per million mapped reads) values of methanol-specific (*mtaCB*), TMA-specific (*mttCB*), DMA-specific (*mtbCB*), and MMA-specific (*mtmCB*) MT1 isoforms in the parent strain (WWM60; orange) and the ΔMA2561 mutant (DDN116; green) during growth on media containing either methanol or TMA as indicated. Significant differences in expression between triplicate cultures of WWM60 and DDN116 using unpaired *t*-tests with Welch correction are shown (*P<0.05, **P<0.01, ***P<0.001, ****P<0.0001), and error bars represent one standard deviation (SD). Red arrows indicate the expression of *mttCB*, *mtbCB*, *mtmCB* genes in the ΔMA2561 mutant on methanol.

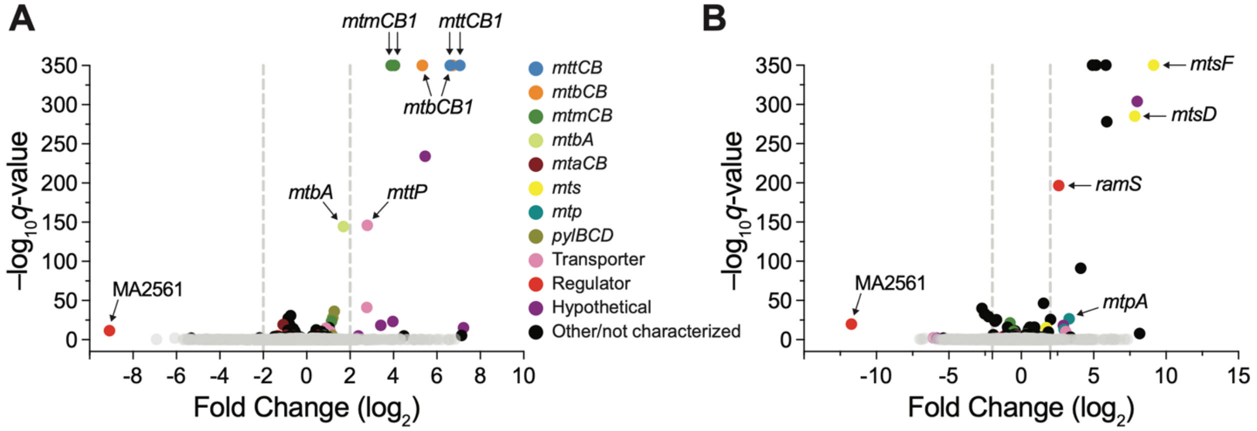


**Figure S14.** Methylamine-specific MT1s are among the most differentially expressed genes in the ΔMA2561 mutant (related to Figure 2D). Volcano plots showing differentially expressed protein-coding genes in DDN116 (ΔMA2561 mutant) compared to the parent strain (WWM60) in media containing either (**A**) methanol, or (**B**) TMA as the sole methanogenesis substrate. Genes without any significant [$-\log_{10}(q\text{-value})<2$] fold change values are shown in gray, while genes with highly significant fold change values ($q$-values below $10^{-310}$) were arbitrarily assigned a $-\log_{10}(q\text{-value})$ of 350 for visualization in the volcano plot.

Gene annotations: *mttCB*: trimethylamine (TMA)-specific methyltransferase complex 1 (MT1); *mtbCB*: dimethylamine (DMA)-specific MT1; *mtmCB*: monomethylamine (MMA)-specific MT1; *mtbA*: methylamine-specific methyltransferase complex 2 (MT2); *mtaCB*: methanol-specific MT1; *mts*: methylsulfide-specific methyltransferase; *mtp*: methylmercaptopropionate-specific methyltransferase; *pylBCD*: pyrrolysine biosynthetic machinery; *mttP*: putative methylamine transporter; *ramS*: homolog of the methylamine methyltransferase corrinoid protein reductive activase.

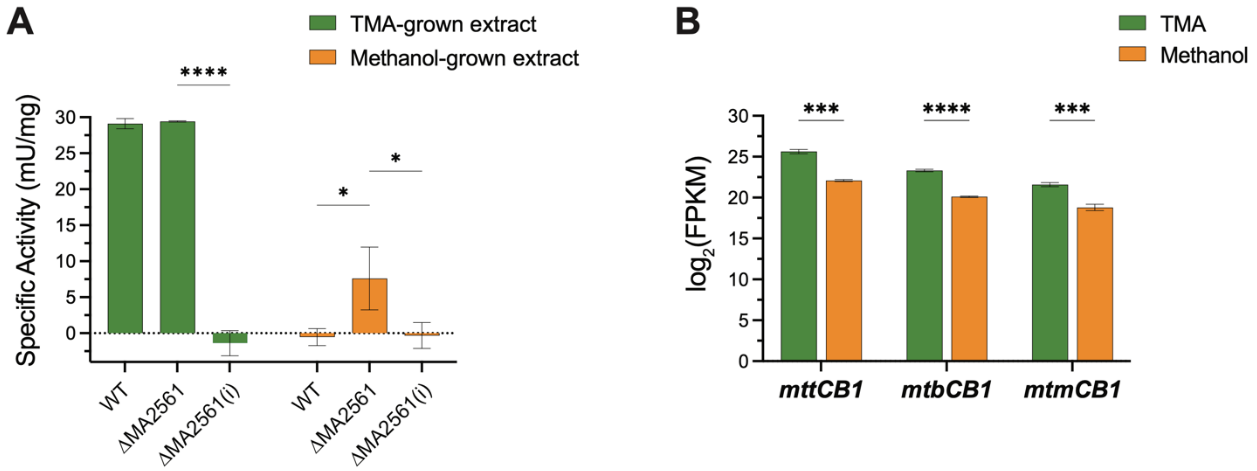


**Figure S15.** Elevated TMA methyltransferase activity and expression in the ΔMA2561 mutant growing on methanol (related to Figure 2E). (**A**) TMA-methyltransferase activity was measured with anaerobic lysates of cells grown in minimal media with either TMA (green) or methanol (orange) as substrate. Specific activity was measured for the parent strain (WWM60), ΔMA2561 extracts, or heat-inactivated ΔMA2561 extracts (i) in triplicate, where 1 U is defined as the amount of enzyme required to consume 1 µmol of coenzyme M per minute. (**B**) Expression level (FPKM) of TMA-, DMA-, and MMA-specific MT1s (*mttCB1*, *mtbCB1*, and *mtmCB1*, respectively) in the ΔMA2561 mutant grown on TMA (green) or methanol (orange). Statistical analyses were conducted using unpaired *t*-tests with Welch correction, where *P<0.05, **P<0.01, ***P<0.001, ****P<0.0001, and error bars represent one standard deviation (SD).

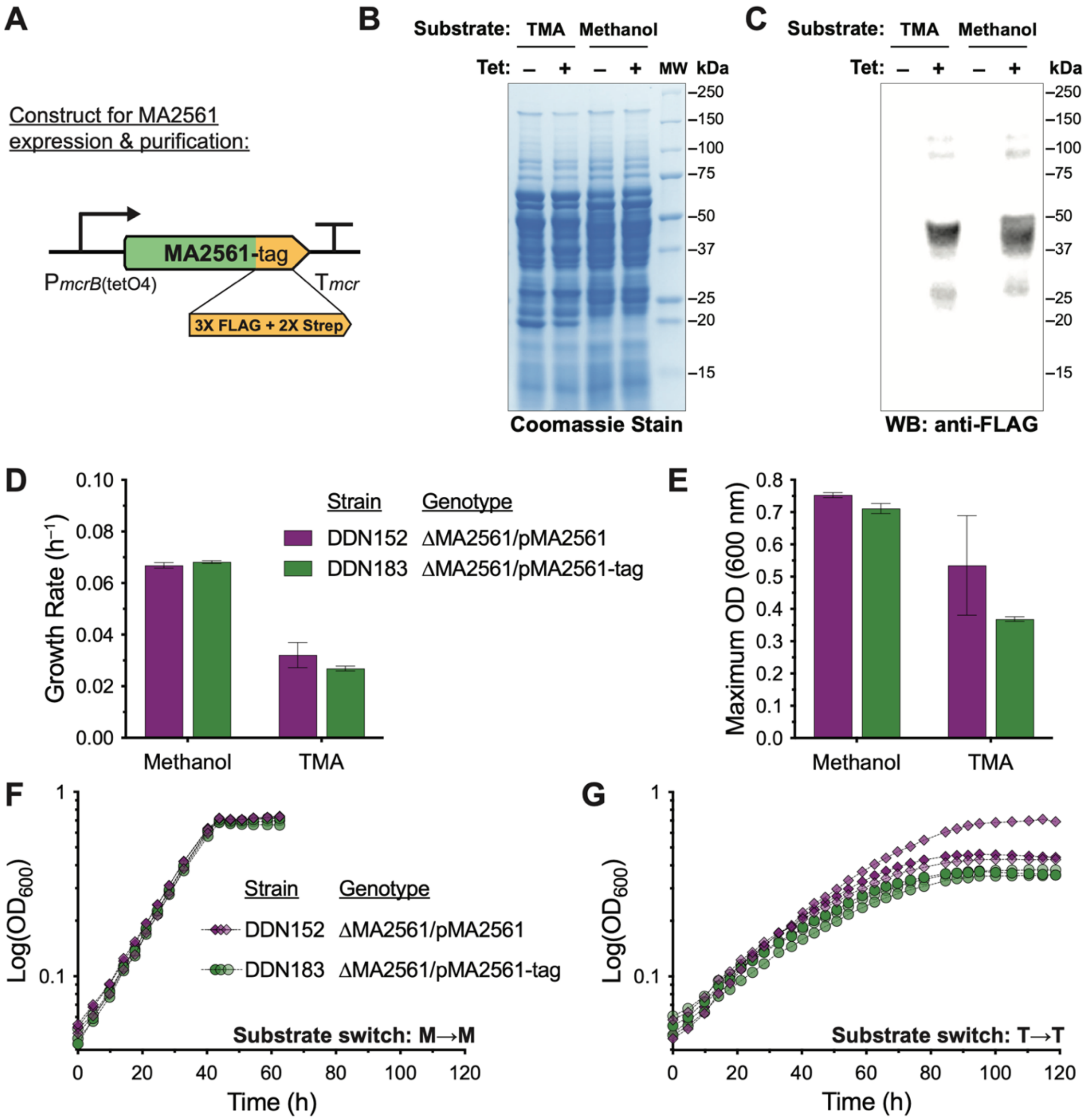


**Figure S16.** Fusion of a C-terminal affinity purification tag does not affect the function of MA2561 *in vivo*. (**A**) A schematic of the vector used to express MA2561 CDS fused to a C-terminal 3X FLAG and 2X Strep tag under the control of the tetracycline-inducible promoter P*mcrB*(tetO4). (**B**) SDS-PAGE of crude cell extracts from DDN183 (the ΔMA2561 mutant with the vector shown in **A**) grown on methanol or trimethylamine (TMA) with (+) or without (–) 100 µg/mL tetracycline as indicated. (**C**) Western Blot using anti-FLAG antibodies to verify expression of the tagged MA2561 locus (expected molecular weight: 45 kDa) with (+) or without (–) 100 µg/mL tetracycline as indicated above the blot. (**D**) Mean growth rate and (**E**) yield from triplicate growth curves of DDN152 (purple) and DDN183 (green) in media with methanol or TMA as the growth substrate and supplemented with 100 µg/mL tetracycline to induce expression. No significant differences were found using unpaired *t*-tests with Welch correction between strains expressing the tagged and untagged MA2561 in each condition. Triplicate growth curves of DDN152 (purple) and DDN183 (green) in media with 100 µg/mL tetracycline on either (**F**) methanol or (**G**) TMA are shown.

**A**

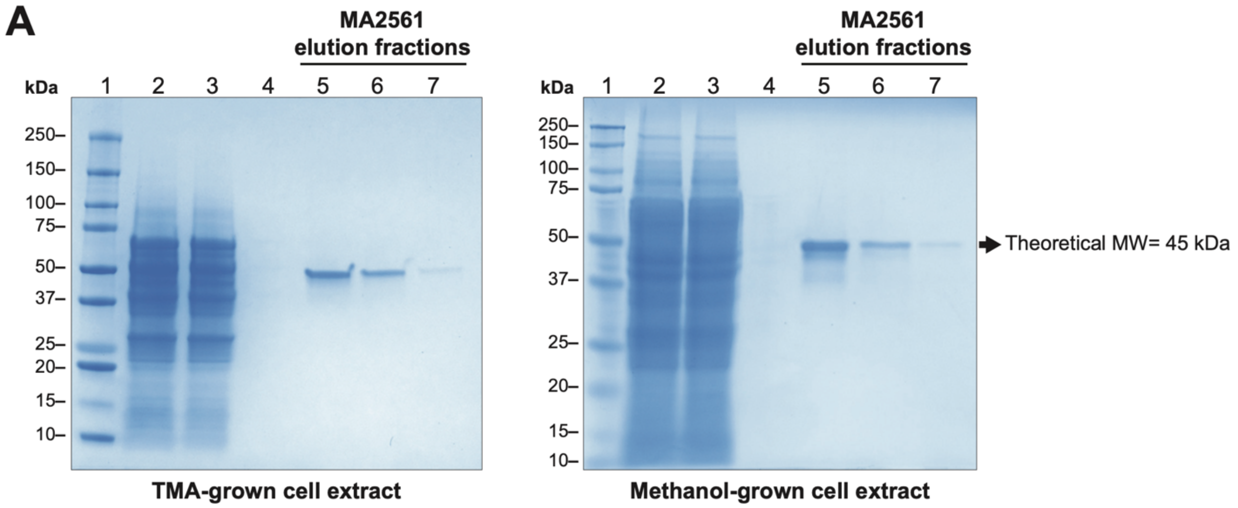


**B**

MAKATSNSTKNGSISKQKACKYLNEMALCSALEMAK**ELISVINKVPVTVFLWRPEKYWPAEFVSEN**
**VKQFGYTVEEFTSGKLLYGNIVHPDDLERVER**ELSR**RIEEDYVDFSQEYR**ILTKSGEVRWVDERTF
IEADENGVVKYLK**GIILDITER**KRK**EKLLYIQR**DLGISLSTSQHLDETLDILLDSCLQIDEIDAGG
IYLVDEDTGDMTLAIQR**GFSPTFVENASYYGANSPNTKLVMIGQPVYKQHIDLLLTSRDDALRQEN**
**LRATAIIPVK**SENEVIAAFYLASSMEYELSDSVR**TVIETIATQFGVFISRIRLEER**LKECVKKRKS

**Figure S17.** Purification of C-terminal tagged MA2561 yields a single ~50 kDa band by reducing SDS-PAGE analysis. (**A**) SDS-PAGE of cell extracts and Strep-TACTIN affinity-purified fractions from *M. acetivorans* DDN183 (ΔMA2561 expressing the C-terminal tagged MA2561; see Figure S16) grown on TMA (left panel, 4-20% acrylamide/bisacrylamide gel) or methanol (right panel, 12% acrylamide/bisacrylamide gel) supplemented with 100 µg/mL tetracycline to induce expression.
Lanes correspond to:
(1) molecular weight standard,
(2) cell extract,
(3) flow-through from column,
(4) the last wash of the column,
(5) first elution fraction,
(6) second elution fraction,
(7) third elution fraction containing a purified protein with an apparent molecular weight of 50 kDa (hypothetical molecular weight of 45 kDa).
(**B**) Amino acid sequence of MA2561 highlighting the region (in red) covered by 19 tryptic peptides identified by mass spectrometry of the 50 kDa-band corresponding to MA2561. Underlined regions show peptides confirmed by fragmentation with an ion score confidence above 99.9% (see Table S6 for MS details).

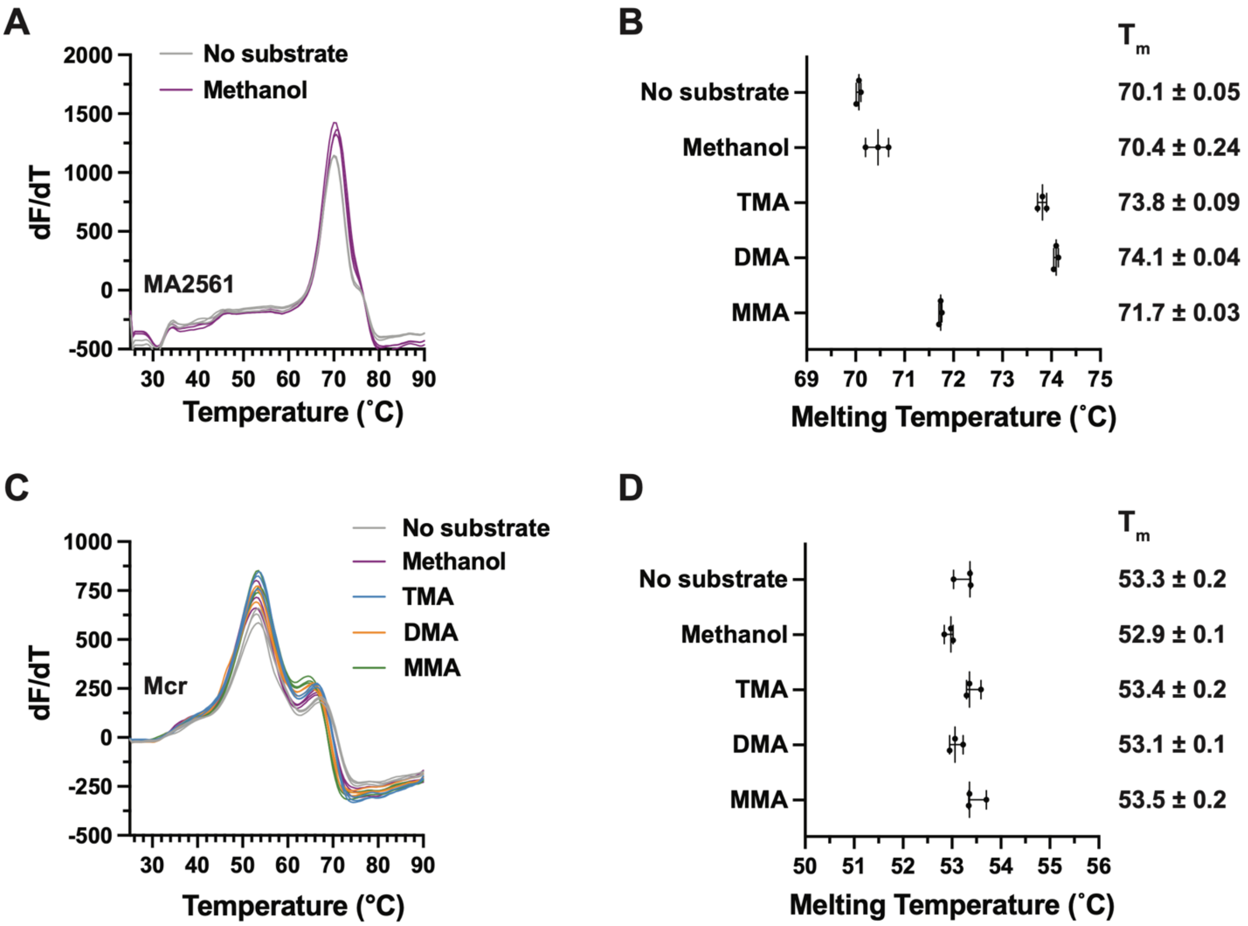


**Figure S18.** Differential Scanning Fluorimetry of MA2561 shows increased thermal stability with methylamines but not methanol (related to Figure 3A). (**A**) Differential Scanning Fluorimetry of MA2561 in the presence (purple) or absence (gray) of 50 mM methanol. (**B**) Box plot of the melting temperature ($T_m$) of MA2561 shows an increase by 2–4 °C in the presence of methylamines (P<0.0001 by unpaired Welch *t*-test). Analysis is based on data from three technical replicates. (**C**) Differential Scanning Fluorimetry of purified Methyl-Coenzyme M Reductase (Mcr) from *M. acetivorans*[2] shows no significant differences in its melting temperature when incubated with 50 mM methanol (purple), 50 mM TMA (blue), 50 mM DMA (orange), or 50 mM MMA (green). (**D**) Box plot of the $T_m$ of Mcr from triplicate melt curves per condition.

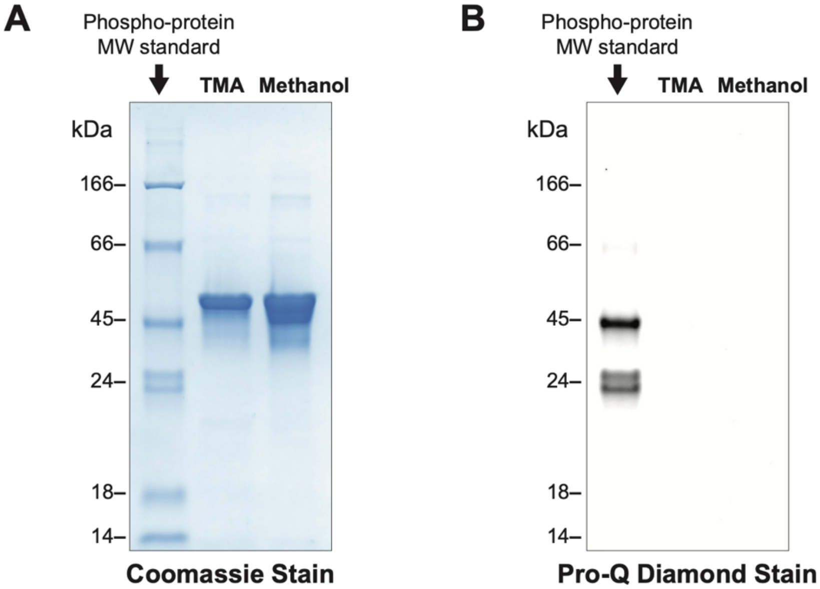


**Figure S19.** MA2561 is not phosphorylated *in vivo*. (**A**) SDS-PAGE analysis (12% acrylamide/bisacrylamide) of Strep-TACTIN-purified and concentrated MA2561 from *M. acetivorans* DDN183 cultures (ΔMA2561 expressing the C-terminal tagged MA2561; see Figure S16) grown on TMA or methanol supplemented with 100 µg/mL tetracycline (to induce protein expression) show no signs of phosphorylation by (**B**) Pro-Q Diamond phosphoprotein staining. The molecular weight standard (Thermo peppermint stick) shows phosphoproteins of 45 and 24 kDa that serve as a positive control for the Pro-Q Diamond phosphoprotein stain.

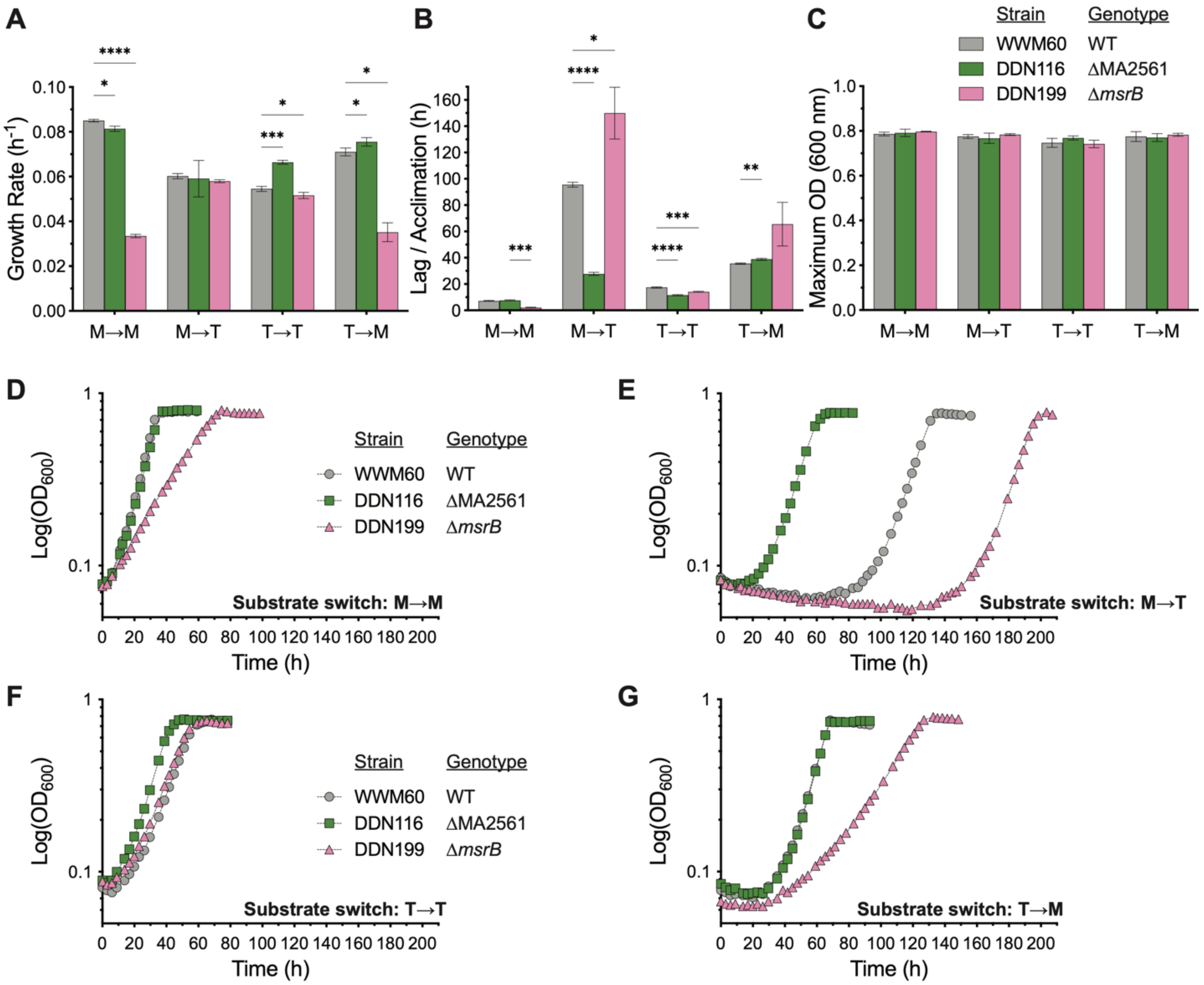


**Figure S20.** ΔMA2561 and Δ*msrB* mutants do not phenocopy each other. (**A**) Growth rate, (**B**) lag/acclimation time, and (**C**) growth yield obtained from triplicate growth curves of the parent strain (WWM60; gray), ΔMA2561 mutant (DDN116; green), and Δ*msrB* mutant (DDN199; pink) when either methanol grown cells are transferred to methanol (M→M) or trimethylamine (M→T), or TMA-grown cells are transferred to TMA (T→T) or methanol (T→M). Statistical analyses of comparisons to the parent strain (WWM60) were conducted using unpaired *t*-tests with Welch correction, where *P<0.05, **P<0.01, ***P<0.001, ****P<0.0001, and error bars represent one standard deviation (SD).

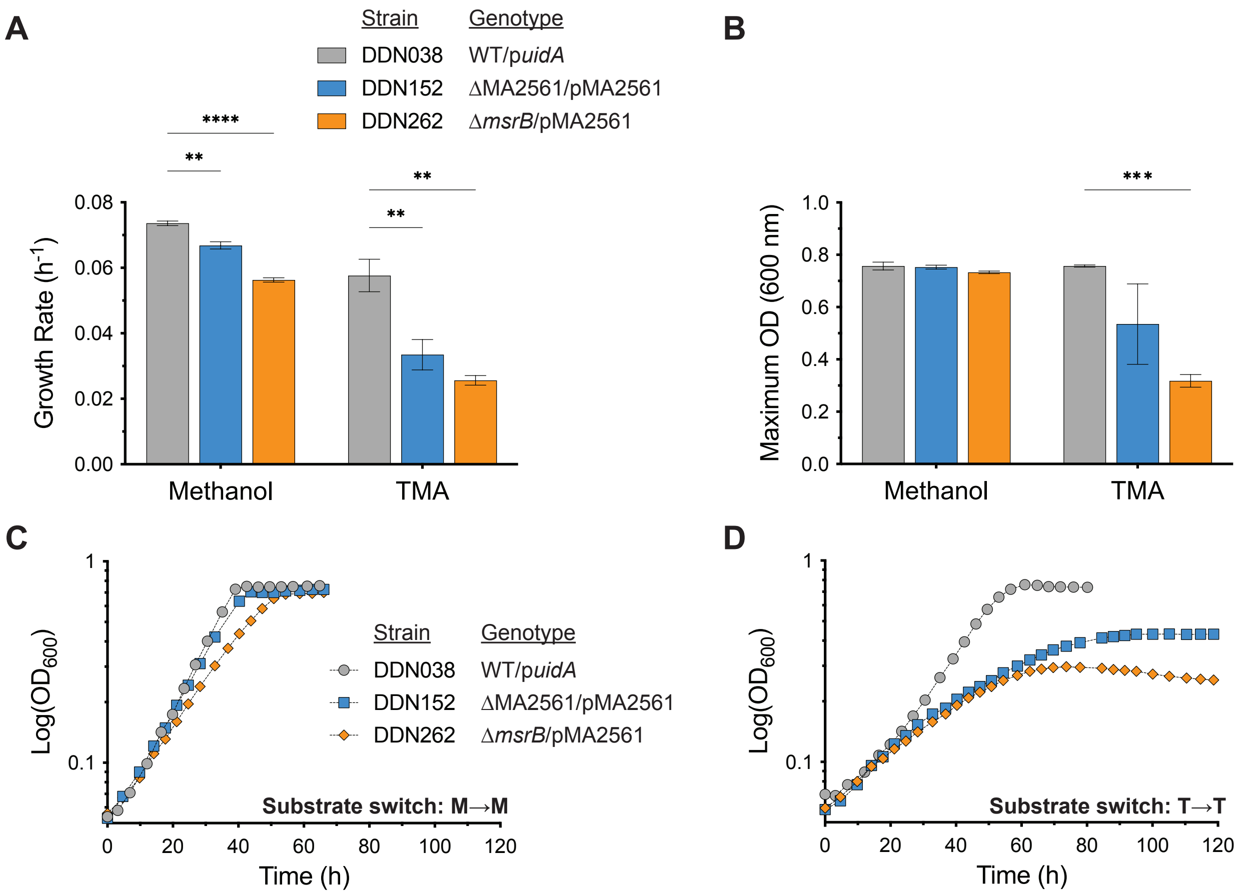


**Figure S21.** Epistasis analysis shows no genetic interaction between MA2561 and *msrB in vivo*. (**A**) Growth rate and (**B**) growth yield from triplicate growth curves on methanol or TMA of the parent strain carrying a plasmid with the *E. coli* β-glucuronidase gene as a negative control ('p*uidA*') (DDN038; gray), ΔMA2561/pMA2561 (DDN152; blue), and Δ*msrB*/pMA2561 (DDN262; orange). DDN152 and DDN262 contain an expression vector with MA2561 ('pMA2561') under the control of the tetracycline-inducible promoter P*mcrB*(tetO4). All growth assays were conducted in media supplemented with 100 µg/mL tetracycline to induce protein expression. Statistical analyses of comparisons to DDN038 were conducted using unpaired *t*-tests with Welch correction, where *P<0.05, **P<0.01, ***P<0.001, ****$P$<0.0001, and error bars represent one standard deviation (SD). Representative growth curves for DDN038 (gray), DDN152 (blue), and DDN262 (orange) in minimal media with (**C**) methanol or (**D**) TMA are shown.

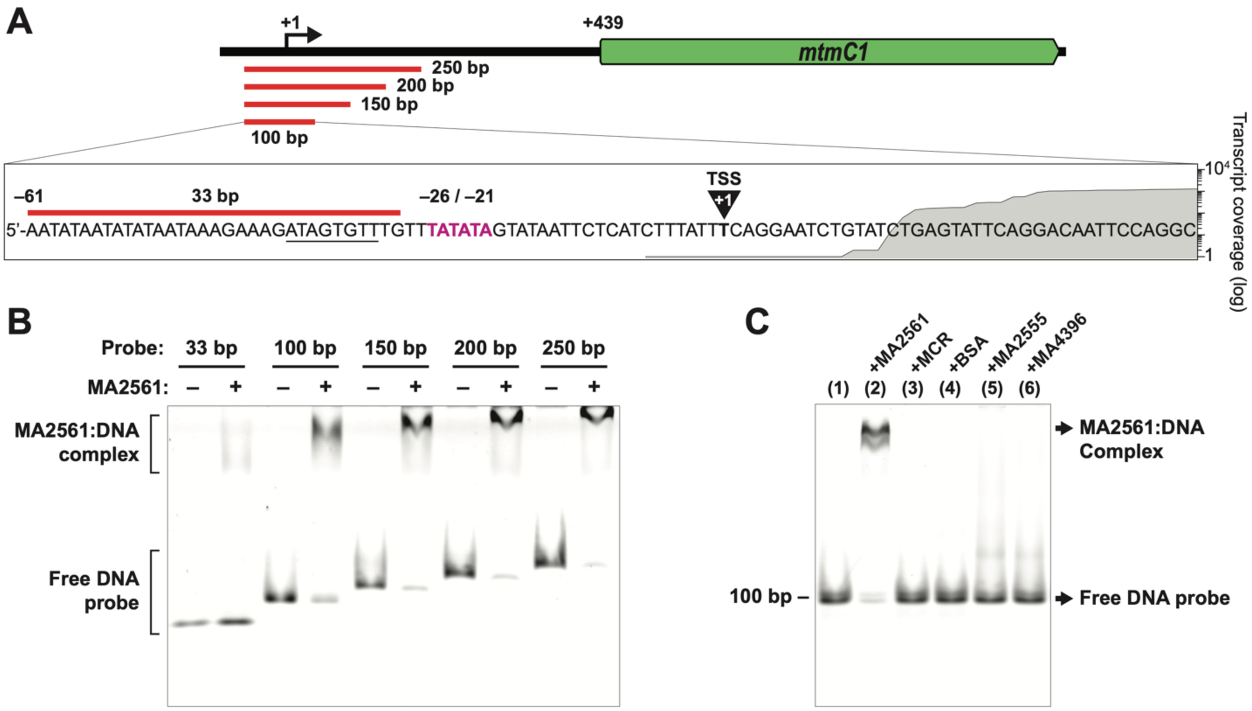


**Figure S22.** MA2561 binds the promoter of the *mtmCB1* operon (related to Figure 3B). (**A**) A map of the 5'-untranslated region (UTR) upstream of the operon encoding the monomethylamine (MMA)-specific MT1 gene isoform 1 (*mtmCB1*). In red, the five different Cy5-labeled probes spanning a 250 bp region used in this study are shown. The inset contains the 100-bp probe sequence highlighting the TATA box (purple) and the transcription start site (TSS) according to Burke *et al*. (1998)[3]. The underlined sequence upstream of the TATA box is the potential B-recognition element (BRE). The transcript coverage of triplicate cultures of WWM60 during growth on TMA are also mapped to this region. (**B**) Electrophoretic Mobility Shift Assay (EMSA) of the DNA probes spanning the *mtmCB1* promoter shown in (**A**), with or without 0.7 mg/mL MA2561. (**C**) EMSA of the 100-bp probe (lane 1) in the presence of 1.3 mg/mL MA2561 (lane 2), 1.3 mg/mL methyl-coenzyme M reductase (Mcr, lane 3), 1.3 mg/mL BSA (lane 4), 1.3 mg/mL MA2555 (lane 5), or 1.3 mg/mL MA4396 (lane 6).

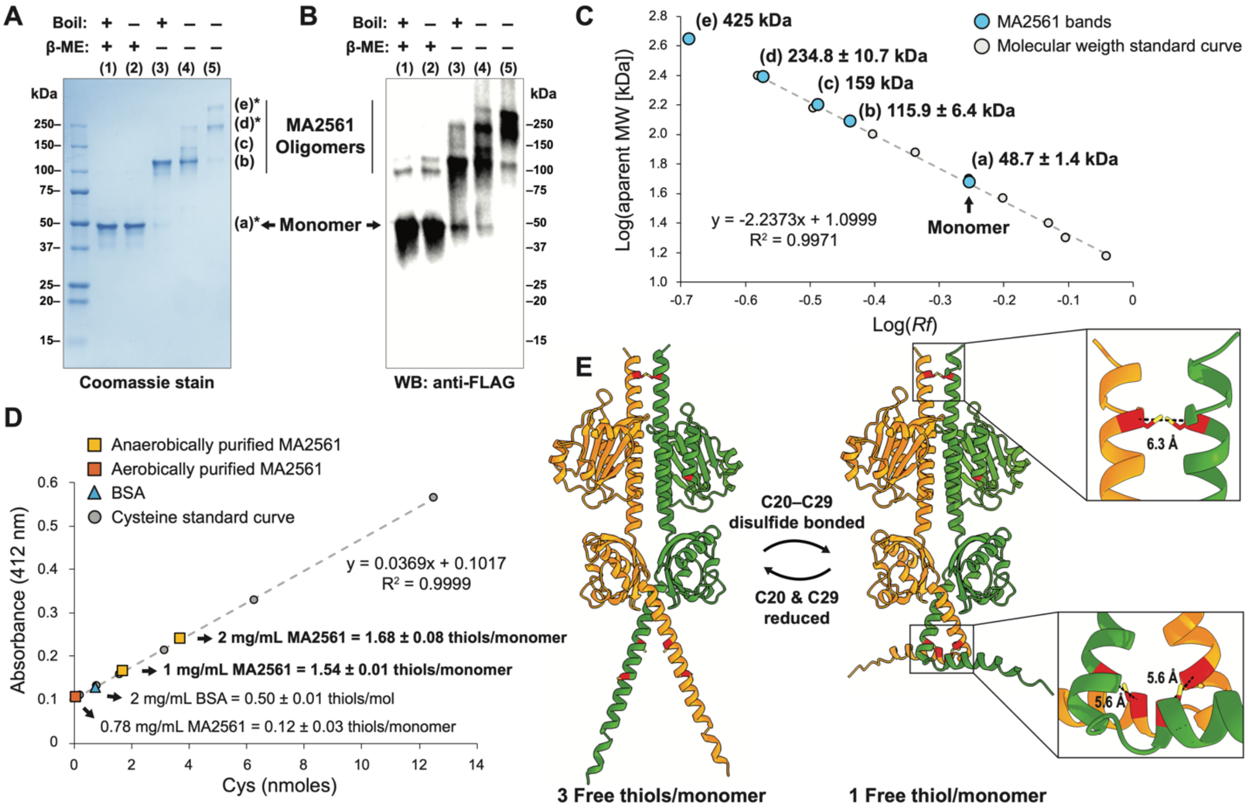


**Figure S23.** MA2561 forms oligomers that are stabilized by interchain disulfide bonds (related to Figure 3E). (**A**) SDS-PAGE and (**B**) anti-FLAG Western Blot of anaerobically (lanes 1–4) or aerobically (lane 5) purified MA2561 treated with or without a reducing agent (+/– β-mercaptoethanol, β-ME), with or without a boil step (5 mins at 100 ºC), prior to loading as indicated. Lanes 1–4 were purified anaerobically, while lane 5 corresponds to an aerobic preparation of MA2561. Distinct and immunoreactive bands corresponding to MA2561 are indicated (a–e), and bands additionally confirmed by mass spectrometry (Table S6) are highlighted with an asterisk. The hypothetical molecular weight (MW) of the tagged MA2561 monomer is 45 kDa. (**C**) Apparent MW calculations of the MA2561 bands shown in (**A**) from triplicate gels plotted against the MW standard curve. (**D**) Quantification of the average oxidation state of cysteines per monomer from two individual anaerobic preparations of MA2561 (in bold, yellow squares) and one aerobic preparation of MA2561 (orange square). Free thiols were quantified using Ellman's reagent and plotted against a standard curve generated with different concentrations of cysteine. Results obtained for BSA, which contains a free exposed cysteine, are shown as a control. MA2561 likely is a homodimer covalently tethered by 1–3 disulfide bonds. (**E**) Alphafold[4] predictions of MA2561 dimers for two conformations depending on the oxidation state of the cysteines at positions 20 and 29. Individual polypeptide chains are shown in the same color, and cysteine residues are highlighted in red (sulfur atom in yellow). Insets show details of potential C20–C29 and C324–C324 interchain disulfide bonds, indicating the distance between the alpha carbons of these residues.

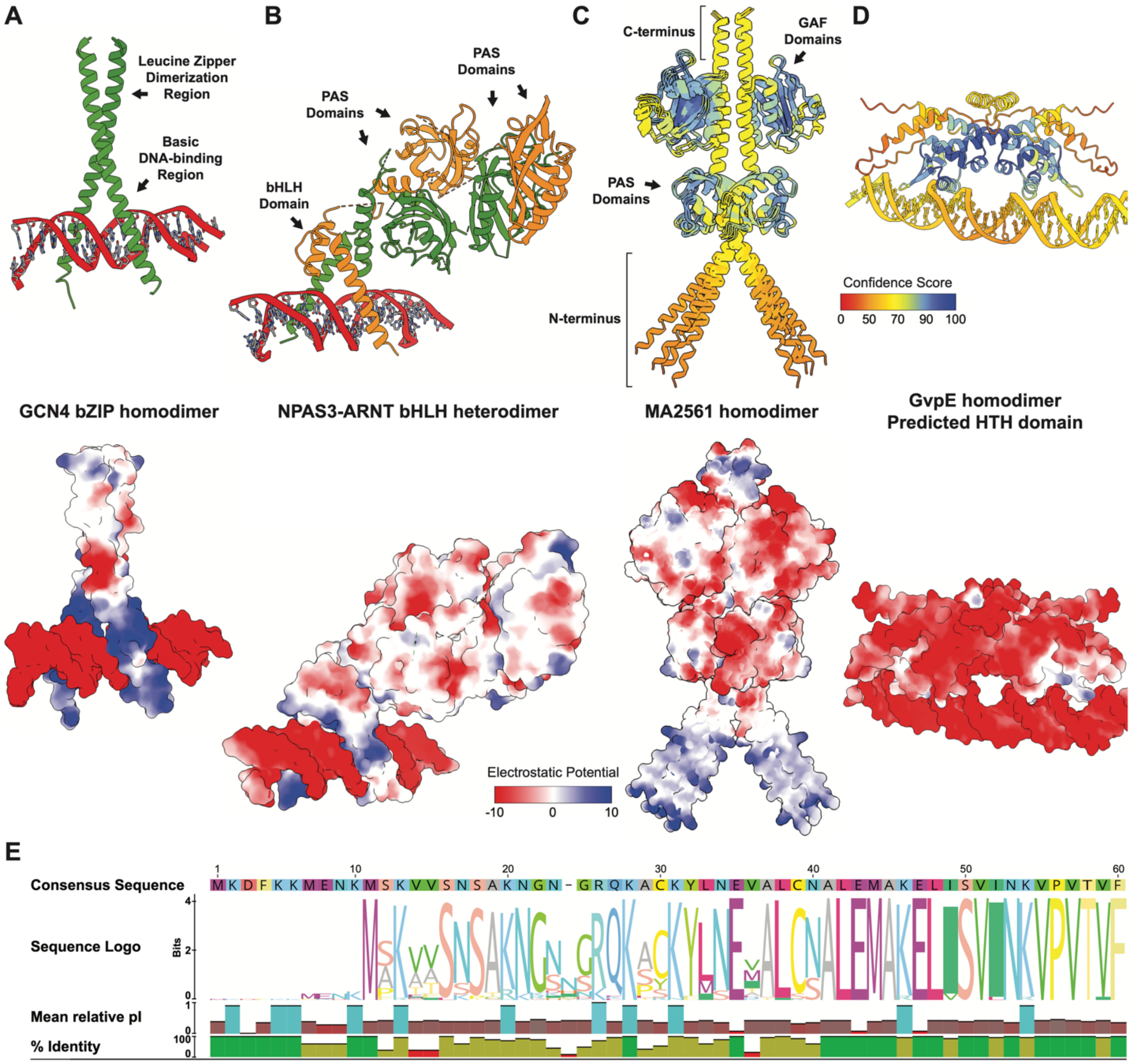


**Figure S24.** MA2561 dimer has a positively charged coiled-coil motif that resembles the basic leucine zipper (bZIP) and basic helix-loop-helix (bHLH) DNA-binding domains found in eukaryotic transcription factors (related to Figure 3F). (**A**) Crystal structure of the yeast bZIP transcription factor GCN4 binding DNA as a parallel left-handed coiled-coil homodimer (top). Surface charge colored by Coulombic electrostatic potential (bottom) shows the basic (blue) region of the GCN4 leucine zipper gripping the DNA major groove. Structure of the bZIP-DNA complex based on Ellenberger *et al*., 1992[5] (PDB ID: 1YSA). (**B**) Crystal structure of the mouse Neuronal PAS Domain-containing Protein 3 and Aryl Hydrocarbon Receptor Nuclear Translocator (NPAS3-ARNT) heterodimer featuring a bHLH domain complex with DNA, in addition to two pairs of PAS domains towards the C-terminus of the dimer (top, ribbon diagram colored by the polypeptide chain). Like bZIP domains, the positive charge of bHLH domains counteracts the negatively charged DNA to form the protein-DNA complex (bottom, surface electrostatic potential). Structure based on Wu *et al*., 2016[6] (PDB ID: 5SY7). (**C**) Superimposed Alphafold models of four dimeric MA2561 conformers highlights a highly flexible N-terminal domain with low predicted local distance difference test (pLDDT) scores, suggesting a poorly structured region or a domain that only acquires a stable conformation as part of a complex (top, ribbon diagram colored by the Alphafold pLDDT per-residue score of local confidence). The flexile N-terminal region also displays a parallel coiled-coil-like motif with a net positive charge (bottom, colored by electrostatic potential) that strongly resembles the scissor-like structure of bZIP and bHLH domains required for DNA binding. (**D**) Alphafold model of a GvpE dimer from the archaeon *Halobacterium salinarum* (NCBI Reference Sequence: WP_010904106.1; top, ribbon diagram; bottom, surface electrostatic potential) and 35 base pairs containing the GvpE-responsive element sequence. Although GvpE was previously proposed to contain a bZIP domain[7,8], its structural model predicts a helix-turn-helix domain of the ArsR family (InterPro 102.0[1]) that is common in bacterial and archaeal transcription factors. (**E**) Sequence logo of the N-terminal region for 62 homologs of MA2561 from isolates within the Genus *Methanosarcina*, indicating the percent identity and the relative isoelectric point of the conserved basic residues.

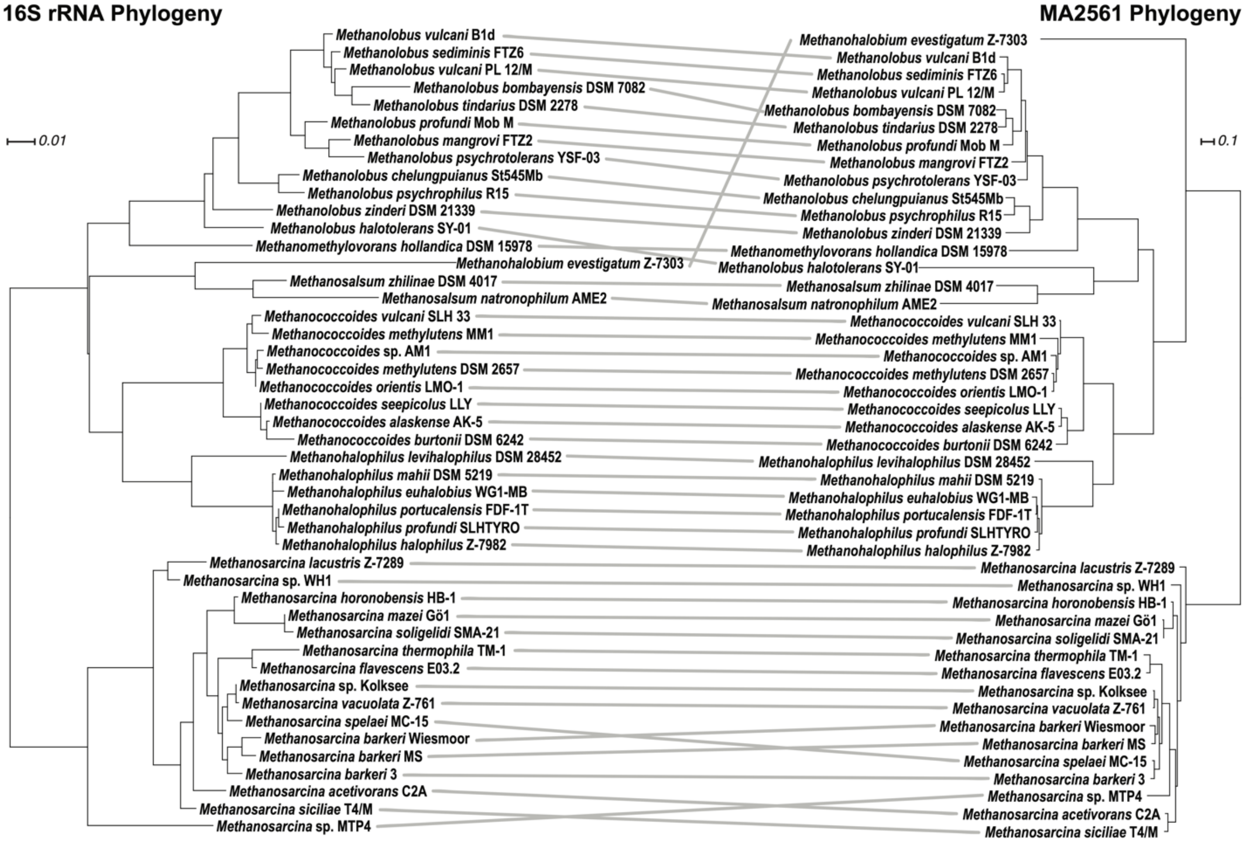


**Figure S25.** MA2561 is vertically inherited in methylotrophic methanogens within the Family *Methanosarcinaceae* (related to Figure 4A). A tanglegram of the maximum likelihood phylogenetic tree of the 16S rRNA gene (left) and the maximum likelihood phylogenetic tree of the amino acid sequence of MA2561 (right) from 46 methanogens that have at least one copy with 40% or more sequence homology to MA2561. Only one MA2561 homolog per strain that is most closely related to *M. acetivorans* MA2561 was used in this analysis.

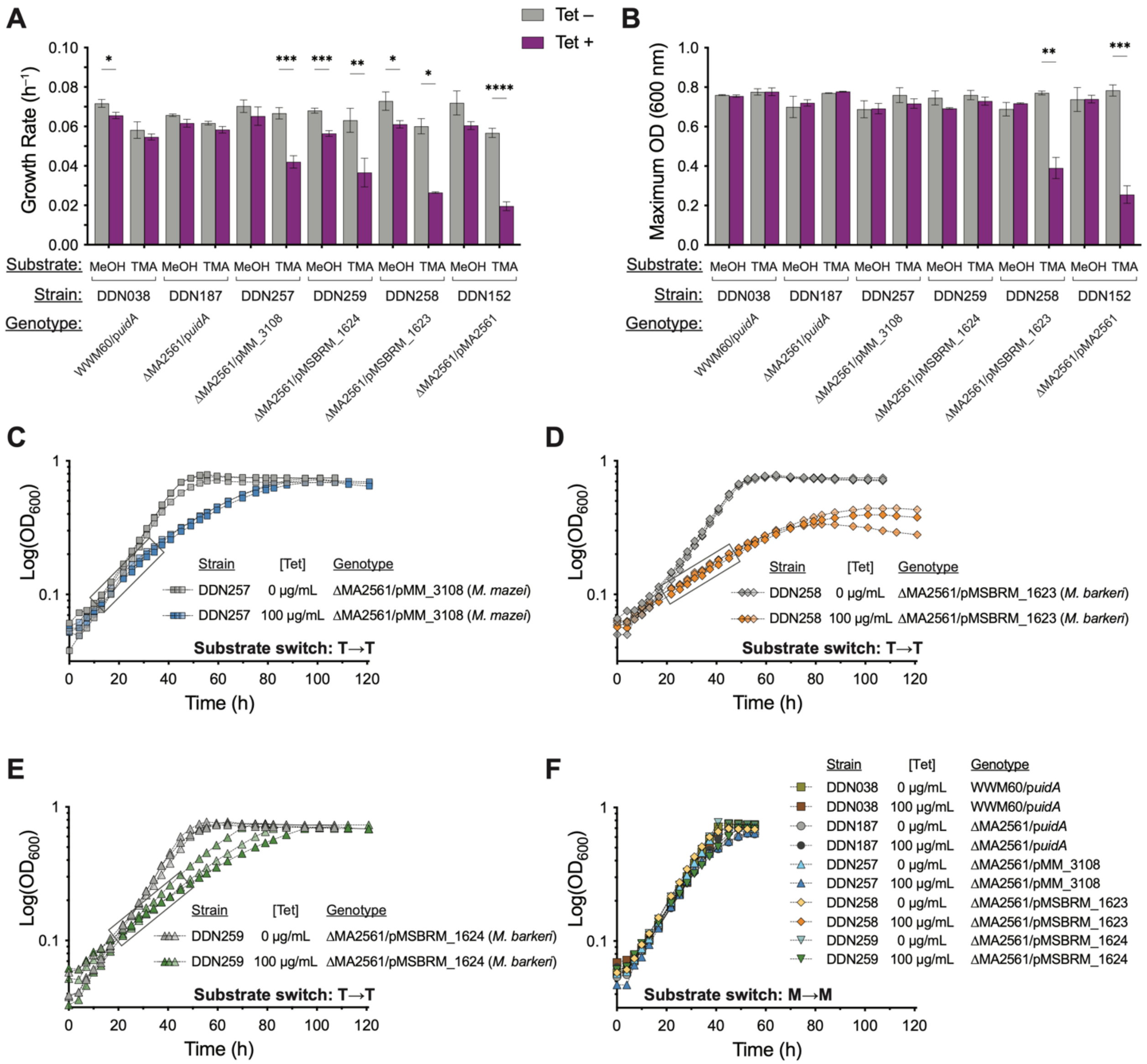


**Figure S26.** The cellular function of MA2561 is widely conserved (related to Figure 4C). (**A**) Growth rate and (**B**) yield from triplicate growth curves of the ΔMA2561 mutant expressing MA2561 homologs from other methanogens *in trans* on an expression vector under the control of the tetracycline inducible promoter P*mcrB*(tetO4). DDN257 contains a plasmid expressing MM_3108 from *Methanosarcina mazei* Gö1, DDN258 and DDN259 contain plasmids expressing MSBRM_1623 and MSBRM_1624 from *M. barkeri* MS, respectively. Control experiments were conducted with the parent strain (WWM60) and the ΔMA2561 mutant expressing the *E. coli* β-glucuronidase gene *uidA*. Data for growth on either methanol or trimethylamine (TMA) are shown as indicated either in the absence of tetracycline (no induction; gray) or in the presence of 100 μg/mL tetracycline (full induction; purple). Statistical analyses were conducted using unpaired *t*-tests with Welch correction, where *P<0.05, **P<0.01, ***P<0.001, ****P<0.0001, and error bars represent one standard deviation (SD). Triplicate growth curves in media with TMA (**C**) for DDN257 (gray—no tetracycline addition; blue—100 μg/mL tetracycline addition), (**D**) DDN258 (gray—no tetracycline addition; orange—100 μg/mL tetracycline addition), and (**E**) DDN259 (gray—no tetracycline addition; green—100 μg/mL tetracycline addition). Data points used to calculate the growth rate in cells overexpressing MA2561 homologs on TMA are enclosed in a rectangle. Data from panels C, D, and E are summarized in Figure 4C and show here for comparison. (**F**) Representative growth curves of strains expressing MA2561 homologs from different *Methanosarcina* strains on methanol with or without tetracycline supplementation as indicated. No substantial or significant difference in growth on methanol was observed across all conditions tested.

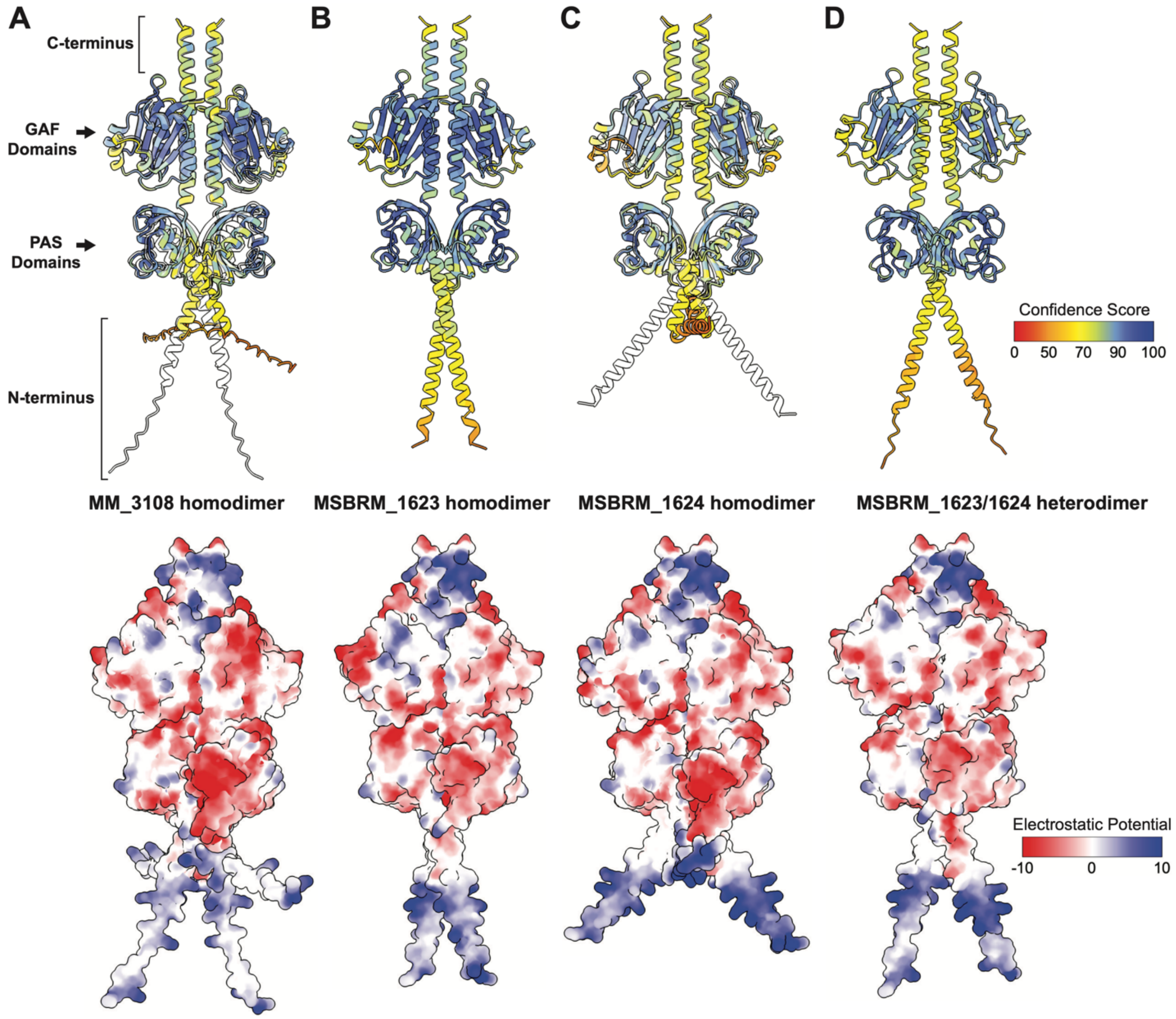


**Figure S27.** Predicted structures of MA2561 homologs preserve a putative bZIP-like DNA-binding motif. Alphafold models of (**A**) *M. mazei* Gö1 MM_3108, (**B**) *M. barkeri* MS MSBRM_1623 and (**C**) MSBRM_1624 homodimers, and (**D**) MSBRM_1623/1624 heterodimer. Ribbon diagrams (top panels) in A and C feature a potential disulfide bond that bends the N-terminal α-helices. Reduction of cysteine, simulated by C20S and C19S mutations for MM_3108 and MSBRM_1624, respectively (superimposed models in white), straightens the N-terminal α-helices. Electrostatic surface potential (bottom panels) shows the presence of a basic N-terminal motif similar to bZIP and bHLH DNA-binding domains.
